# Unraveling the intracellular cross-talk governing the balance between TNFα mediated survival and apoptosis signaling

**DOI:** 10.1101/2021.09.07.459274

**Authors:** Sharmila Biswas, Baishakhi Tikader, Sandip Kar, Ganesh Viswanathan

## Abstract

Tumor necrosis factor alpha (TNFα), a pleiotropic cytokine, helps maintain a balance between proliferation and apoptosis in normal cells. This balance is often sacrificed in a diseased cell, such as that of a cancer, by preferring survival phenotype over apoptosis. Restoring this balance requires a detailed understanding of the causal intracellular mechanisms that govern TNFα stimulated apoptotic response. In this study, we use a systems biology approach to unravel the interplay between the intracellular signaling markers that orchestrate apoptosis levels. Our approach deciphered the synergism between the early intracellular markers phosphorylated JNK (pJNK) and phosphorylated AKT (pAKT) that modulate the activation of Caspase3, an important apoptotic regulator. We demonstrate that this synergism depends critically on the survival pathway signaling mediated by NFκB which plays a dominant role in controlling the extent of the overall apoptotic response. By systematic inhibition of the signaling markers, we establish that the dynamic cross-talk between the pJNK and pAKT transients directs the apoptosis phenotype via accumulated Caspase3 response. Interestingly, superposition of the semi-quantitative correlation between apoptosis and Caspase3 transient levels on the proposed TNFα network model permits quantification of the dynamic apoptotic response under different stimulation conditions. Thus, the predictive model can be leveraged towards arriving at useful insights that can identify potential targeted therapeutic strategies for altering apoptotic response.

## Introduction

The pleiotropic cytokine Tumor necrosis factor alpha (TNFα) mediates diverse cellular phenotypic decisions such as apoptosis, inflammation, proliferation (Aggarwal, 2003; Hoffmann and Baltimore, 2006; Locksley et al., 2001; Mantovani et al., 2008; Micheau and Tschopp, 2003; Newton and Dixit, 2012; Sankar and Michael, 2002; Wallach et al., 1999; Zelová and Hošek, 2013). Under various conditions, large quantities of TNFα is secreted by tissue-resident cells such as monocytes, macrophages, T-lymphocytes in tissue-microenvironment (Beutler et al., 1985; Ha et al., 2008; van Horssen et al., 2006; Mercogliano et al., 2021). In normal cells, TNFα maintains a balance between different phenotypes (Aggarwal et al., 2012). Since such a balance is disrupted in a diseased cell in a cancer or autoimmune milieu (Jarosz-Griffiths et al., 2019; Kearney et al., 2018; Lopez and Tait, 2015; Sherekar and Viswanathan, 2021; Shishodia and Aggarwal, 2002), its restoration using interventional therapeutic approach involving TNFα is being considered recently (Kaufman and Choi, 1999; Vilcek, 2008). TNFα is used as a drug in a variety of cancer (Aggarwal, 2003; Balkwill, 2009; van Horssen et al., 2006; Kay and Rahman, 2009; Kreuzaler and Watson, 2012; Sharma et al., 2011; Wu, 2009) and immune-related therapies (Kodama et al., 2005; Steinman et al., 2012), in novel chemotherapy strategies (Holoch and Griffith, 2009; Lu et al., 2012; Rana et al., 2009) and in combination therapies (some of which are currently in different phases of clinical trials) (Amm et al., 2011; Gregorc et al., 2011; Herman et al., 2013; Pilati et al., 2008). Therefore, insights into how TNFα orchestrates different phenotypic response, particularly that of apoptosis is paramount to improving the existing therapeutic approaches and arriving at novel ones.

Evading apoptosis, a form of regulated cell-death, is one of the hallmarks of cancer (Fouad and Aanei, 2017; Hanahan and Weinberg, 2000, 2011; Lopez and Tait, 2015). TNFα signaling and the associated intracellular entities control mechanisms that regulate cell-death (Janes et al., 2005; Kist and Vucic, 2021; Vince et al., 2007). One of the primary events involved in cells exhibiting an apoptotic response is activation of Caspase3. Caspase3 is one of the effector caspases and is a key player involved in regulation of cell-death (Kist and Vucic, 2021; Reed et al., 1996; Shishodia and Aggarwal, 2002; Wajant et al., 2003). Cells when exposed to TNFα triggers a cascade of signal transducing proteins such as MAPKs, Bcl2 placed appropriately in the underlying wiring diagram and thereby facilitates direct and indirect regulation of activated Caspase3 (Duffey et al., 2000; Galluzzi et al., 2018; Kist and Vucic, 2021; Takada and Aggarwal, 2004; Yang et al., 2016). Along with modulating cell-death, TNFα strongly controls the proliferation as well as survival responses by transducing relevant information through entities such as NFκB (Van Antwerp et al., 1996; Charles et al., 2009; Liu et al., 2017; Reed et al., 1996; Werner et al., 2008), phosphorylated AKT (pAKT) which is activated by PI3K (Burow et al., 2000; Kandel et al., 2002; Meyer et al., 2012; Vivanco and Sawyers, 2002). Even though these key regulators have been identified long before to be involved in proliferation and apoptosis (Gerondakis et al., 1999; Meffert et al., 2003), it is as yet unclear how these entities synergistically regulate the overall long-term apoptotic outcome. A detailed understanding of the dynamics of each of these entities and their inter-dependence in the context of apoptotic regulation is necessary to distill out the overall response by a certain cell-type stimulated by TNFα.

Molecular network (Behar and Hoffmann, 2010; Nurse, 2008; Purvis and Lahav, 2013) that regulates a cell’s response to TNFα would consist of several entities and interactions between them that facilitates information exchange (Aggarwal, 2003; Chen and Goeddel, 2002; Schleich and Lavrik, 2013). Network dynamics is dictated by the context-specific nature of the activation of these entities and the associated transient properties (Adlung et al., 2017; Deng et al., 2003; Dhanasekaran and Reddy, 2008; Guo et al., 1998; Lamb et al., 2003; Meyer et al., 2012; Ventura et al., 2006; Zhang et al., 2004). While TNFα driven transient phosphorylated JNK (pJNK) activation enhances cell survival, its sustained activity promotes cell death by inhibiting growth signals via ERK (Leaner et al., 2003; Ventura et al., 2006). On the other hand, absence of activation of pAKT due to blocking of PI3K results in attenuating its inhibitory effect on Caspase3 leading to a delayed induction of cell-death (Burow et al., 2000). Moreover, a combined computational and perturbative experimental study showed that along with other intracellular phospho-proteins, pAKT may be involved in influencing the stimulus-strength dependent kinetics of the short-term (4 h) TNFα signaling induced apoptotic response (Gierut et al., 2015; Lau et al., 2012). These studies suggest that in shorter time scales, pAKT could be a key modulator of TNFα induced apoptosis via Caspase3 activation. However, apoptosis in mammalian systems occurs over much larger timescales. This demands investigation of the TNFα triggered Caspase3 activation and its regulation by the intracellular proteins over a sustained period of time. In TNFα signaling network, these intracellular proteins are interlocked via feedback (Sabio and Davis, 2014) and feedforward (Leaner et al., 2003; Oliver Metzig et al., 2020) loops, and other direct and indirect pathways facilitating transient information exchange between them when activated (Aggarwal, 2003; Deppamann and Janes, 2013). In these loops and pathways, NFκB acts as a key mediator (Bouwmeester et al., 2004, Liu et al., 2017).

NFκB transients in a cell stimulated by TNFα contain sufficient information to sense the stimulus dose and accordingly fine-tune the regulation of cellular responses (Zhang et al., 2017). Recent study on NIH3T3 cells demonstrated that NFκB dynamics precisely captures the rate of change in dose of TNFα stimulation (Son et al., 2021). Single-cell level modelling of the TNFα dose-dependent NFκB signaling revealed that a cell’s commitment to apoptotic or proliferative phenotype could be embedded in the early transient response of NFκB (Lipniacki et al., 2007). NFκB regulates several entities that inhibit various effector caspases, including Caspase3 via multiple pathways and thereby exhibits a strong influence on the apoptotic response due to TNFα (Shishodia and Aggarwal, 2002; Tang et al., 2019). This indicates that when NFκB signal is arrested, Caspase3 level should increase making TNFα treated cells culminate in apoptosis. Since Caspase3 is also independently activated by TNFα via Caspase8 (Micheau and Tschopp, 2003; Wang et al., 2008), a question arises as to what is the extent of synergism that is brought in by arresting NFκB downstream signaling.

In order to assess the nature of the above highlighted synergism, we hypothesized that dynamic evolution of the key entities such as pJNK and pAKT along with the transient activity of NFκB on Caspase3 could play a key role in the regulation of apoptosis in response to TNFα. Herein, the goal of this study is to unravel this dynamic interaction between the key intracellular markers that regulates Caspase3 downstream and subsequently predict apoptotic response. We took an experimental data-constrained systems biology approach to decipher the dynamic correlation of the intracellular state and the overall phenotypic apoptotic outcome as a response to TNFα.

## Results

### 1. NFκB signal inhibition enhances TNFα mediated apoptotic response

To study the dynamics of cellular fate on TNFα stimulation, we measured the apoptosis fraction using Annexin V assay (Methods) in the U937 model cell line. A 24 h exposure of TNFα (100 ng/ml) results in 25% of the population undergoing cell-death (Figure 1A, blue). Low cell-death following exposure to only TNFα indicates that in U937 cells the cytokine might be favoring the survival signaling as well (Aggarwal, 2003) and maintains a balance between the apoptotic and survival responses. Since NFκB is a strong mediator of survival signaling, TNFα likely stimulates NFκB signaling in U937 cell lines too. In order to ascertain the involvement of NFκB, we inhibited NFκB signaling by exposing cells with Triptolide (TPL), a known inhibitor of NFκB which blocks its transactivation and thereby blocking the signal flow of the survival pathways (Lee et al., 1999). Pre-treatment with TPL (60 nM) and subsequent exposure to TNFα resulted in 80% cell death (Figure 1A, black). A comparison of apoptosis levels for this case with those corresponding to only TPL (Figure 1A, red) pre-treatment determines that blocking survival pathway in U937 can lead to sufficient apoptotic response. However, those cells experiencing a pre-treatment of TPL showed a greater initial (8 h) apoptotic response in the presence of TNFα (Figure 1A, red and black). At the same time, the levels of cell death in the case of only TPL, and those pre-treated with TPL and exposed to TNFα were similar at 24h. This indicates that the survival signaling pathway coordinates with those of apoptosis in order to regulate the cell-death in U937 cells. However, the exact control mechanism governing this coordination remains unclear.

**Figure 1.**
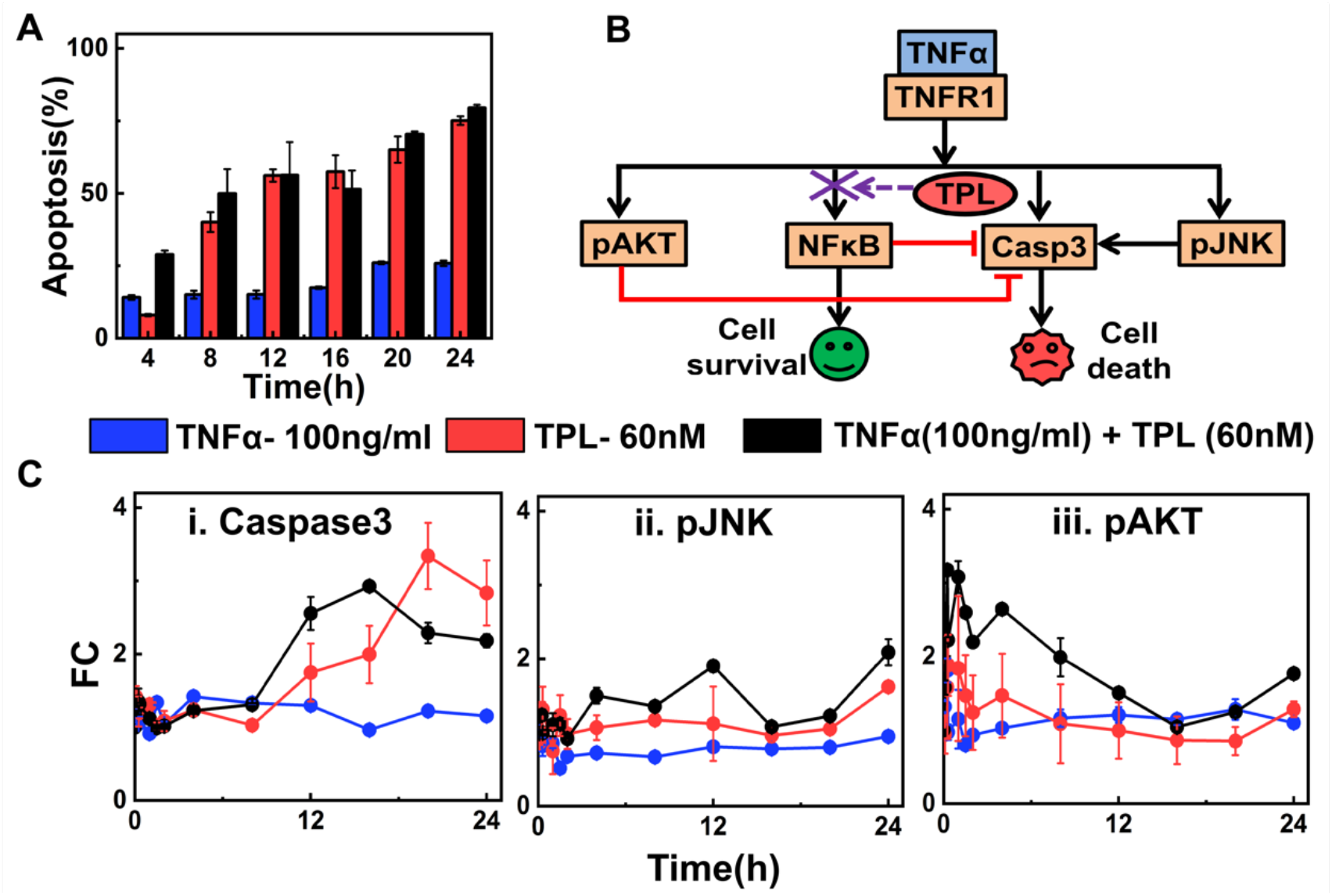
Apoptosis and intracellular signaling patterns due to TNFα stimulation in the presence and absence of NFκB activity. (A) Cell death percentage measured using Annexin V assay under three different experimental conditions at different time points. (B) A schematic of TNFα signaling mediating survival and apoptotic responses via key intracellular regulators. (C) Trajectories of relative fold change (FC) of (i) Caspase3, (ii) pJNK, and (iii) pAKT for three different experimental conditions. Experiments were done in triplicates using U937 cells. Details of extraction of apoptosis percentage are in Text S1, Supplementary information.

Deciphering the control mechanism requires distilling out the intracellular dynamic features that orchestrate the survival and apoptotic responses in U937 cells stimulated with TNFα. The key regulatory entities involved in this orchestration are depicted in Figure 1B. It is well established that the two upstream signaling markers pAKT and pJNK regulate apoptosis (Gierut et al., 2015; Leaner et al., 2003; Ventura et al., 2006). pAKT has a significant role in preventing Caspase3 activation (Ozes et al., 2005). While pJNK has a function in activating downstream caspases (Deng et al., 2003), committing cells to apoptosis. Moreover, the role of pJNK has been demonstrated to be context-specific (Deng et al., 2003; Lamb et al., 2003). Thus, we simultaneously measured the dynamic levels (in terms of relative fold change (FC)) of these activated signaling proteins using high-throughput experimentation (Krutzik and Nolan, 2006; Manohar et al., 2019) (Methods; Text S1, Supplementary Information), which are presented in Figure 1C. In the case of only TNFα stimulation, no significant difference in Caspase3 dynamics (Figure 1C(i), blue) was observed which corroborates with the low apoptosis percentage (Figure 1A, blue). Note that even for pJNK and pAKT trajectories for the case of only TNFα stimulation, an initial increase in the fold change was observed (Figure 1C(ii-iii), blue). On the other hand, under NFκB inhibitory conditions (TPL and TNFα+TPL), a rise in the later time period (>8h) of the Caspase3 FC (Figure 1C(i), red and black) commensurates with the enhanced cell death as found in Figure 1A (red and black). The subsequent continuous decrease in pAKT is reflected by sustained increase of Caspase3 levels.

These results suggest that cells undergo enhanced apoptosis with change in the dynamics of the three marker proteins, *viz*., pAKT, pJNK and Caspase3, if NFκB mediated survival signaling is inhibited by using TPL. However, the extent of dynamical crosstalk between the intracellular marker proteins cannot be inferred from such observations. It remains unclear how the dynamics of upstream proteins refine the downstream Caspase3 response, eventually regulating the apoptosis. It remains inconclusive as to which entities could be tuned to regulate the cellular decision-making events involved in cell survival or apoptosis. Thus, to understand how the crosstalk between multiple intracellular dynamic signals quantitatively modulates cellular apoptosis, we need to develop a predictive mathematical model of the underlying network.

### 2. Data-driven mathematical model of TNFα network

A wiring diagram of the TNFα network considered for the data-driven mathematical model is depicted in Figure 2A. Biochemical details of the interactions in Figure 2A are presented in Text S2.1 and Figure S3, Supplementary Information. The network model (Figure 2A) assumes that TNFα on binding to TNFR1 activates MAPK cascades (pERK and pJNK), NFκB and pAKT survival pathways along with the Caspase3 pathway, which mediates apoptosis (Aggarwal, 2003). Molecular players such as ceramide are also activated upon TNFα stimulation, which subsequently influences the pAKT and pJNK levels (Sawada et al., 2004). As NFκB inhibition resulted in an augmented level of apoptosis (Figure 1), we included nodes that are governed by NFκB, and are involved in activation or inhibition of the three marker proteins. Key nodes mediating crosstalk between the marker proteins leading to apoptosis regulation were incorporated (Aikin et al., 2004; Wicovsky et al., 2007; Wullaert et al., 2006). Note that while some of the interactions in the network (Figure 2A and Figure S3) are direct activation or inhibition, rest capture the overall causal effect due to a complex set of sequential biological interactions. (Details of these are given in Table S1, Supplementary Information.)

**Figure 2.**
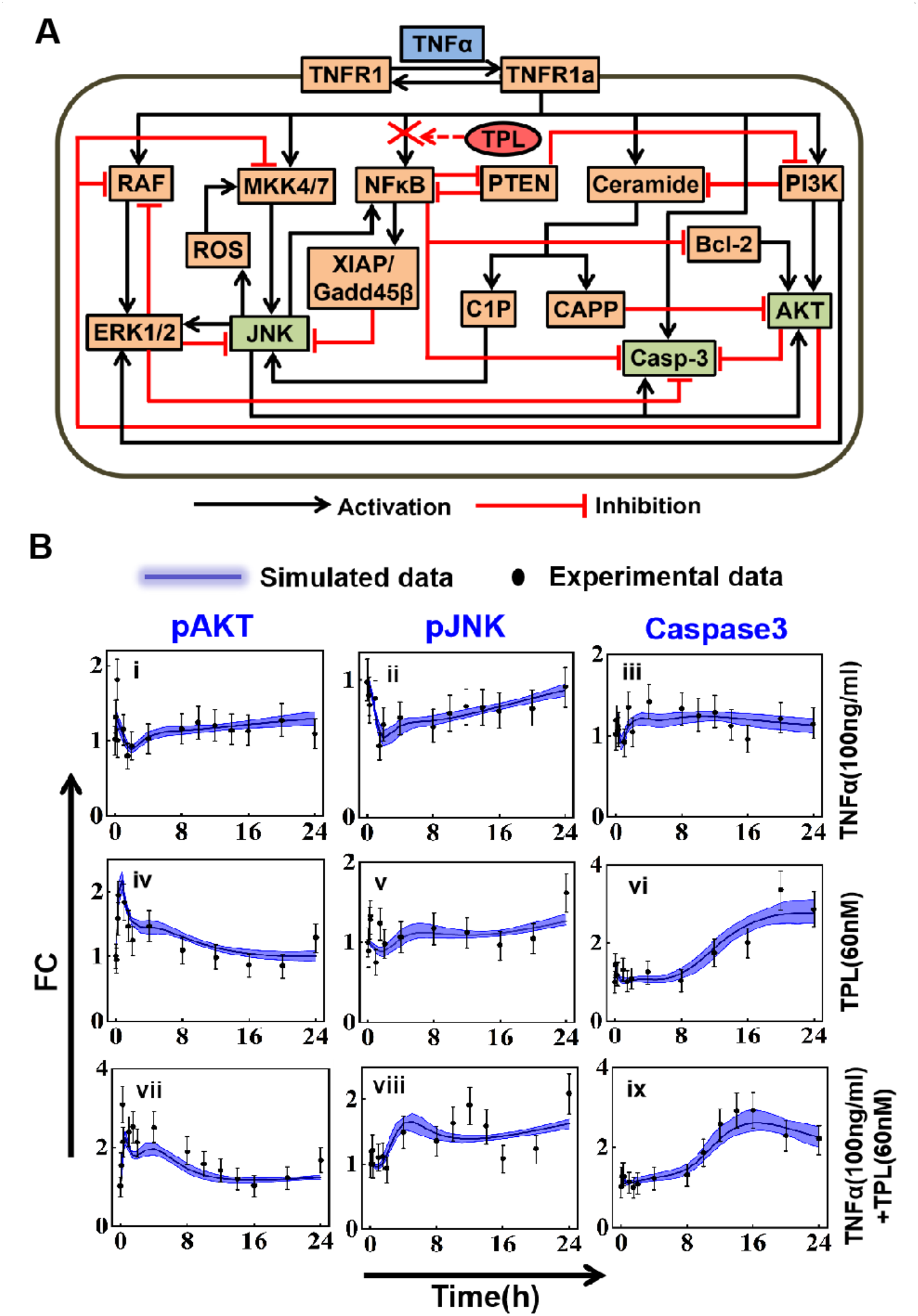
Interaction network and model training with quantitative time resolved data of U937 cells. (A) Schematic of TNFα signaling network. Black arrows and red hammers, respectively represent activation and inhibition of entities. Network is stimulated by TNFα (blue box). The experimentally measured marker proteins are designated green boxes. Inhibition of NFκB by TPL (red oval) is shown by a red cross. TNFα upon binding to TNFR1 leads to activation of pERK and pJNK via RAF and MKK4/7, respectively. MKK4/7, ROS and pJNK are involved in a positive feedback loop. TNFα activates NFκB, PI3K, and Caspase3. Multiple pathways from NFκB inhibiting Caspase3 are lumped as a single inhibitory interaction. Various cross-talks between the entities are incorporated. (Details of the entities and the interactions are in Section 2, Supplementary Information). (B) Comparison of the model simulated transients of pAKT (i, iv, vii), pJNK (ii, v, viii), and Caspase3 (iii, vi, ix) with the corresponding experimental observations for all three stimulation conditions (TNFα (100 ng/ml), TPL (60 nM), and TNFα (100 ng/ml) + TPL (60 nM)). While columns correspond to those for a signaling marker, the rows correspond to the stimulation conditions. The error bars represent standard deviation calculated with a standard error model (Maiwald and Timmer, 2008) for each experimental data point.

In order to construct a quantitative model of the network (Figure 2A and Figure S3), the interactions in the wiring diagram were quantified by either mass-action kinetics or phenomenological terms. (The model equations and the associated information capturing the dynamics of different entities are provided in Text S2.2 and S2.3, Supplementary Information.) The model was then trained and optimized (Methods) with the experimentally observed relative fold change (FC) of the marker proteins (Figure 1C) to get an estimate of the associated parameters listed in Table S4, Supplementary Information. The model predicted state of the network is contrasted with the training dataset in Figure 2B for all the three stimulation conditions (TNFα, TPL, TNFα+TPL). The model transients for the entities were obtained using the 3% best fit, that is, 60 parameter sets which were identified using the appropriate goodness of fit, χ^2^ value and Akaike information criteria (Methods). (Note that these 3% best fit parameters are statistically well constrained and identifiable as shown in Text S2.4, Supplementary Information.) Model estimated best fit parameter sets adequately capture the experimental relative fold change (Figure 2B). The dynamics obtained for the best fit parameter set along the statistical details are shown in Text S2.5, Supplementary Information. Unless otherwise explicitly stated, best fit parameter set will be used for further analysis.

For the TNFα condition, the model predicted transients for all the three marker proteins are highly consistent with the experimental observations (Figure 2B(i-iii)). For the case of TPL only, the consistency in the prediction of transients is well within the experimental error limits (Figure 2B(iv-vi)). On the other hand, for the third stimulation condition (TNFα+TPL), the model trajectories followed the experimental observations for all the time points considered for pAKT and Caspase3 (Figure 2B(vii, ix)). However, the prediction for the case of pJNK matched with all but a few time points (Figure 2B(viii)). Intriguingly, the model has perfectly identified the transient Caspase3 dynamics for all three stimulation conditions (Figure 2B(iii, vi, ix)).

In order to assess the predictability of the model, we next examine the simulated transients with independent experimental observations obtained for a different condition, that is, lower concentration of TPL (10 nM) in the presence and absence of TNFα (Text S3, Supplementary Information). The model predicted the pAKT and Caspase3 dynamics with reasonable accuracy under both the experimental conditions (Figure S6, Supplementary Information). However, the trajectories for the case of pJNK under both these conditions deviated at certain timepoints (Figure S6, Supplementary Information). This is primarily due to the fact that the deviation of the model predictions and the experimental observations were appreciable for pJNK even in the training set (Figure 2B(viii)).

Overall, the mathematical model of the network curated based on literature reliably mimics our experimental findings. However, mere comparison of the dynamics of the experimentally observed transients of the marker proteins do not offer insights into how the overall dynamics is regulated. We hypothesize that such an insight can be obtained if the cross-talk between various entities in the network is unraveled systematically. Unless otherwise stated explicitly, we will use the model with the identified best parameter sets for distilling out the cross-talk in TNFα signaling network.

### 3. Unraveling the dynamic cross-talk between signaling entities in the network

We next focus on unraveling the cross-talk between different proteins to understand the interdependent regulation of pAKT, pJNK and Caspase3 dynamics. In order to achieve this, we dissected the entire dynamics of the pAKT, pJNK and Caspase3 proteins by analyzing the contributions by specific entities affecting them. The extent of influence by these entities on the marker proteins pAKT, pJNK and Caspase3 is quantitatively captured by the corresponding biochemical reaction rate or flux. Note that the instantaneous level of a certain protein is a linear combination of the contributing fluxes with appropriate sign. (Details of flux calculations along with associated expressions are provided in Text S4, Supplementary Information.) In order to assess the relative influence of various entities over the marker proteins, we track the dynamics evolution of these fluxes (Figure S7, Supplementary Information) under all three stimulation conditions (TNFα, TPL, TNFα+TPL).

We present in Figure 3 dynamic evolution of the fluxes of the interactions that contribute significantly to the transient levels of the marker proteins. The reaction flux analysis shows that pAKT dynamics is positively regulated by pJNK and PI3K, while CAPP is a negative regulator of pAKT under only TNFα stimulation condition (Figure 3A(i)). In the initial phase, pAKT was majorly controlled by pJNK (Figure 3A(i)), thus the drop in the pJNK contribution is reflected on the decreasing levels in the initial phase (upto 4 h) of the dynamics of pAKT (Figure 2B(i)). At later time-points, both pJNK and PI3K mediated effects dominate over other fluxes, and overcome the CAPP facilitated inhibition to initiate the late phase activation of the pAKT dynamics (Figure 3A(i) and Figure 2B(i)). Bcl2 protein does not influence pAKT level, as it’s activity is repressed by NFκB (Figure 3A(i)). However, under NFκB inhibitory conditions (TPL and TNFα+TPL), Bcl2 induces a rapid increase in the reaction flux (Figure 3B) during the initial phase (upto 2 h). This results in a sharp increase in the initial dynamics of pAKT (Figure 2B(iv,vii)). A comparison of the contributions to pAKT for only TNFα stimulation (Figure 3A(i)) and those with TPL stimulation (Figure 3B(i), 3C(i)) suggests that pJNK contributions increased relatively as a result of NFκB inhibition. However, PI3K contribution is completely lost, which can be attributed to the absence of NFκB inhibition of PTEN (Figure 3B(i), 3C(i)). This re-balancing of fluxes ensures a sustained maintenance of the pAKT dynamics at the later phase (Figure 2B(iv,vii)). In the case of only TPL, CAPP mediated inhibition of pAKT is insignificant (Figure 3B(i)) as CAPP activation via Ceramide requires TNFα stimulation.

**Figure 3.**
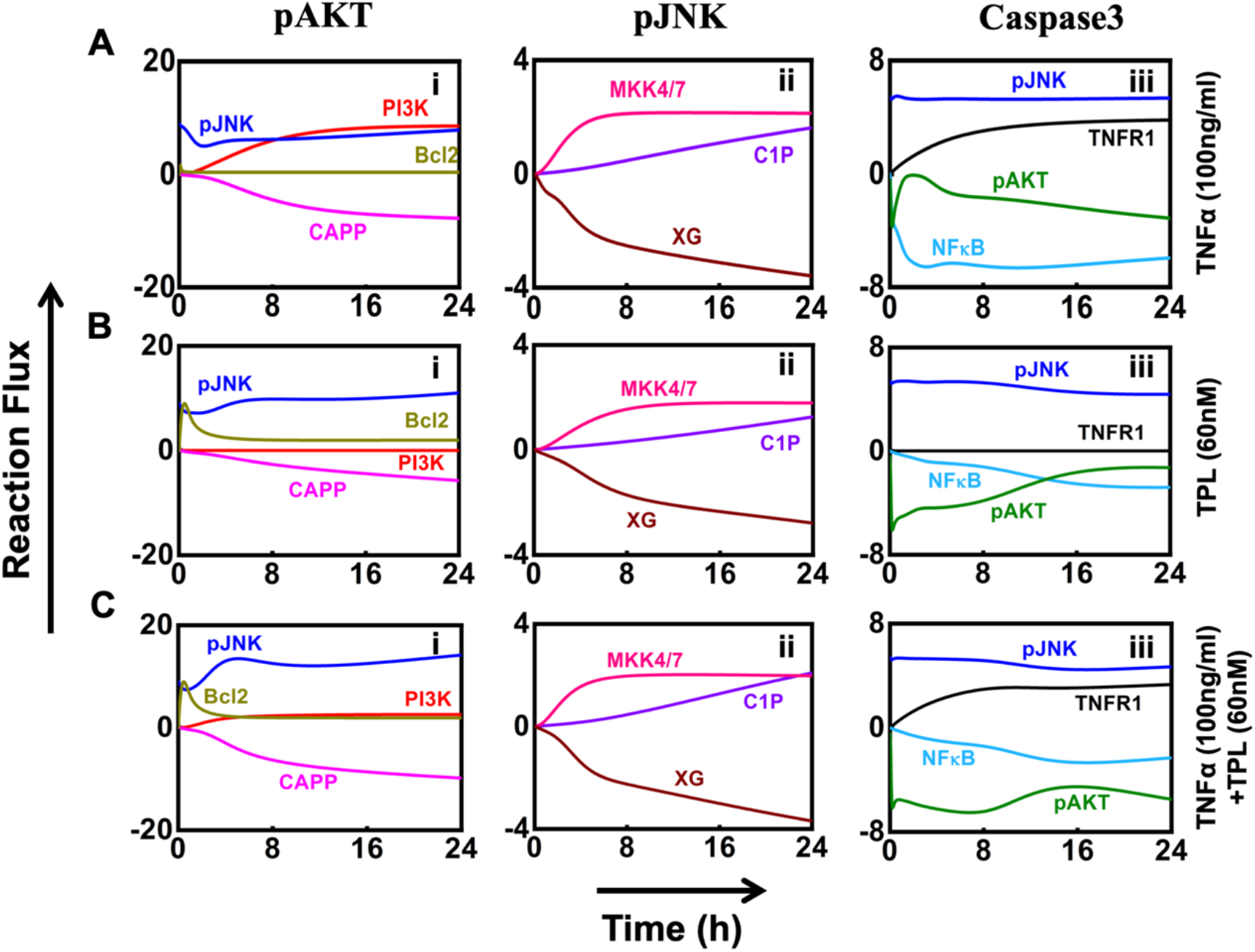
The evolution of majorly contributing fluxes towards pAKT, pJNK and Caspase3. Panels (A), (B), and (C), respectively show the evolution of fluxes for the three markers under the stimulation conditions TNFα, TPL, and TNFα +TPL. In each of the panels (i), (ii), and (iii), respectively show the evolution of fluxes contributing to dynamics of pAKT, pJNK and Caspase3 dynamics for the three different stimulation conditions. The individual reaction fluxes were estimated using the rate equation described in Table S5, Supplementary Information. Evolution of all contributing fluxes is presented in Figure S5, Supplementary Information.

The reaction flux investigation further indicates that C1P and MKK4/7 are the key controllers of pJNK dynamics, while XIAP/Gadd45B (XG) negatively influences it (Figure 3A(ii), 3B(ii), 3C(ii)). Our analysis reveals that initially the inhibitory effect of XG predominates over the contribution from any other nodes under only TNFα stimulation (Figure 3A(ii)), which creates a steep drop in the FC of pJNK (Figure 2B(ii)) in the early phase. In the later phase, the C1P and MKK4/7 primarily control the levels of pJNK (Figure 3A(ii)). However, in the presence of TPL, the inhibition by XG appears to be less effective (Figure 3B(ii), 3C(ii)), which helps to stimulate the dynamics of pJNK at early time points (Figure 2B(v,viii)).

Flux analysis further unfolds that NFκB and pAKT are the major negative regulators of Caspase3 dynamics, while pJNK and TNFR1 are its positive modulators (Figure 3A(iii), 3B(iii), 3C(iii)). When stimulated with only TNFα (Figure 3A(iii)), the balance between the predominant inhibition by NFκB and mutual activation mediated by TNFR1 along with pJNK aid in maintaining sustained levels of Caspase3 (Figure 2B(iii)). Under TPL treatment, the initial dynamics of Caspase3 seems to be influenced both negatively by pAKT and positively by pJNK (Figure 3B(iii)) resulting in no activation of Caspase3 at the early phase (~8h) (Figure 2B(vi)). As the later phase depicts a reduced pAKT inhibitory contribution towards Caspase3 flux (Figure 3B(iii)), there is an augmentation in the level of Caspase3 dynamics (Figure 2B(vi)). Under the influence of TPL in the absence (Figure 3B(iii)) and presence (Figure 3C(iii)) of TNFα, the contribution from NFκB reduced significantly, while pAKT plays a predominant role in maintaining the dynamics of Caspase3 (Figure 2B(ix)). The inhibition mediated by pAKT was relatively less effective and thus controls the threshold level of Caspase3 protein resulting in a sudden increase in the level (Figure S7, Supplementary Information). Note that the positive contribution via pJNK helps in maintaining the sustained levels of Caspase3 under all three stimulation conditions (Figure 3A(iii), 3B(iii), 3C(iii)). However, for TNFα+TPL condition, the negative contribution of pAKT on Caspase3 dynamics starts increasing after 10h (Figure 3C(iii)), leading to a gradual decrease in the Caspase3 levels (Figure 2B(ix)) during the later phase (16-24 h).

In summary, there is a cross-talk between the pAKT and pJNK dynamics which in coordination with NFκB regulates the Caspase3 dynamics. However, our flux analysis does not clearly identify how the Caspase3 transient dynamics is precisely controlled by both pAKT and pJNK under different stimulation conditions. In order assess the nature of modulation exhibited by these two entities on Caspase3 dynamics, we next perform a time-dependent correlation analysis under the three stimulation conditions.

### 4. Dynamic correlation between pJNK and pAKT levels modulates overall Caspase3 transient response

In the previous section, we showed how transient dynamics of pAKT and pJNK individually modulates Caspase3 levels. However, whether this modulation occurs synergistically or not is not evident from the transient dynamics. Since the cooperative influence need not necessarily be instantaneous and could be revealed over an extended period of time, the accumulation of pAKT and pJNK may exhibit synergism in controlling the overall Caspase3 levels along with the apoptotic response. In order to assess this, we perform a systematic time-dependent correlation analysis of the accumulated levels of these markers to decipher the underlying regulation under different stimulation conditions. The accumulation of the marker proteins is quantified by estimating the Area Under the Curve (*AUC*) of their transient response (Methods) over different phases of the dynamics. In particular, we consider accumulation from the start (0 h) of the stimulation up to 8h, 12h and 24h for the correlation analysis. Note that the transients generated using the model for 5 best fit parameter sets were employed for this purpose.

The average relative contributions of accumulated pAKT (*A_pAKT_*) and pJNK (*A_pJNK_*) to overall Caspase3 levels are defined respectively by Eqs (4) and (5) in Methods. The relative contributions for the three stimulated conditions are presented in Figure 4A. When cells were stimulated with just TNFα, pJNK primarily contributes to the Caspase3 accumulated over the various time durations (Figure 4A(i), shaded bars). Under TPL condition, pAKT dominates the Caspase3 accumulation for the first 8 h (Figure 4A(ii), unfilled bars). On the other hand, pJNK, which promotes Caspase3 activation, contributed negatively to Caspase3 accumulation (Figure 4A(ii), shaded bars). Accumulation up to 12 h shifted the pJNK contribution from negative to positive, while pAKT contribution decreased (Figure 4A(ii)). Subsequently, the influence of these two markers on overall Caspase3 level up to 24 h becomes poised (Figure 4A(ii)). In the case of TNFα+TPL, the positive contribution from pJNK towards Caspase3 accumulation over 8, 12 and 24 h increases along with a decrease in the impact of pAKT on it (Figure 4A(iii)). This indicates that pJNK plays a significant role in regulating Caspase3 expression throughout the entire time course under all three stimulations conditions except during the initial phase for only TPL treatment.

**Figure 4.**
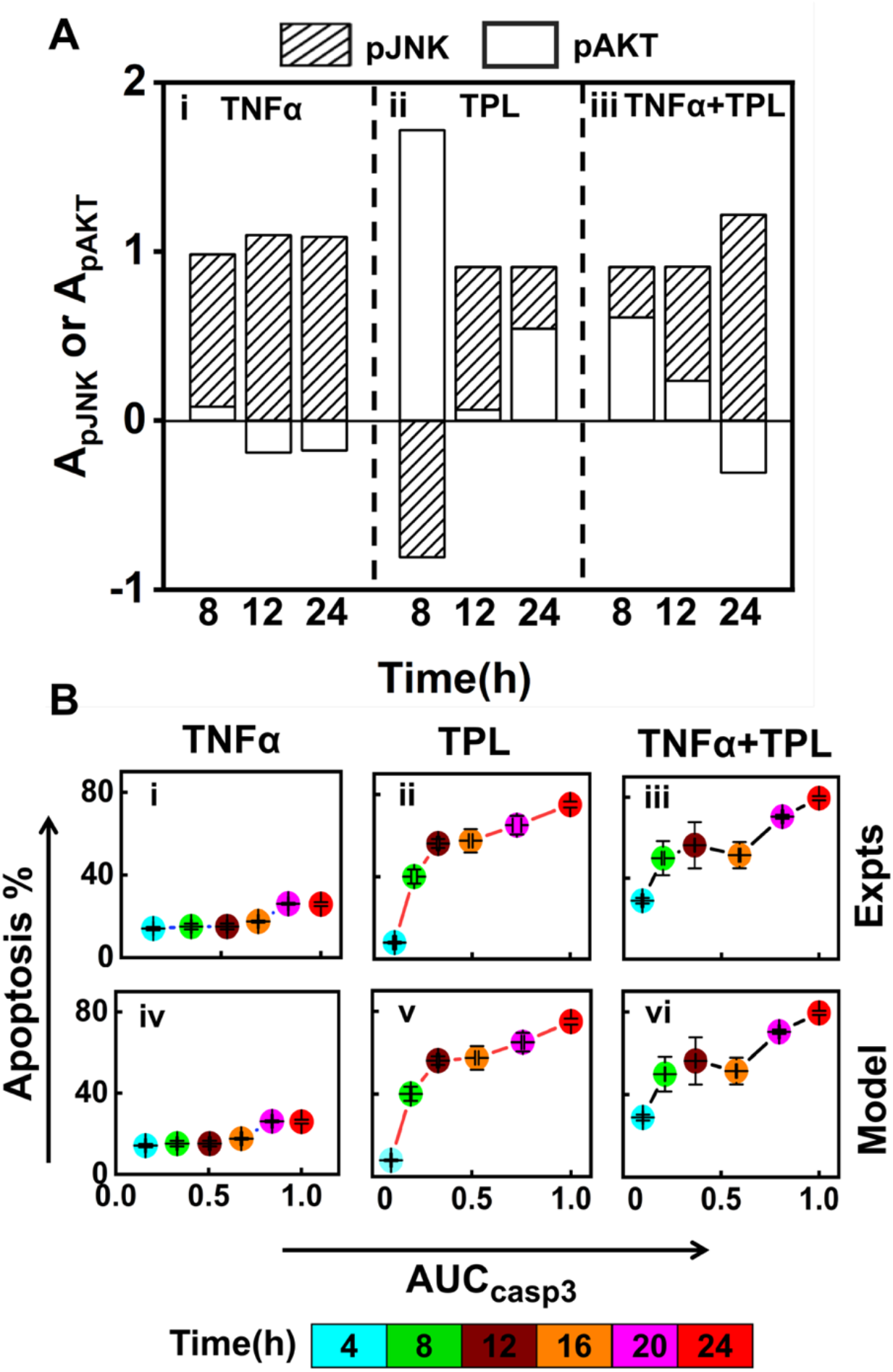
pJNK and pAKT transients synergistically control the overall Caspase3 levels and modulate the apoptotic response. (A) Relative contribution of model generated pJNK and pAKT transients corresponding to 5 best fit parameter sets (*A_pJNK_* and *A_pAKT_*) towards the corresponding overall Caspase3 accumulation for 8, 12 and 24 h durations for (i) TNFα, (ii) TPL, (iii) TNFα+TPL stimulation. (B) Depicts the correlation of experimentally measured accumulated Caspase3 levels (*AUC_casp3_*) (n=3) with corresponding apoptosis percentage for the stimulation conditions (i) TNFα, (ii) TPL, (iii) TNFα+TPL. (iv-vi) represent correlation of model predicted accumulated Caspase3 levels (*AUC_casp3_*) (n=5) with the experimentally measured apoptosis percentage (n=3) for the three stimulation conditions. The time points are appropriately color-coded in (B).

Since Caspase3 along with other effector caspases such as Caspase7 are known to initiate apoptotic response (Wajant et al., 2003), we determine how apoptosis is reflected in the overall Caspase3 accumulation (*AUC_casp3_*) under these three stimulation conditions (Figure 4B). First, we analyze the experimentally measured apoptosis levels (Figure 1A) with the corresponding accumulation of Caspase3 (Figure 1C(iii)). For TNFα stimulation condition, the apoptosis percentage did not rise concomitantly with the increasing *AUC_casp3_* (Figure 4B(i)). This could probably be due to the activation of the survival pathway (NFκB), which interferes with the apoptosis process (Figure 2A). When the NFκB mediated survival signals are interrupted using TPL, there was an enhancement of Caspase3 accumulation over a 24h time period with an increase in the apoptosis level (Figure 4B(ii)). Moreover, the proportion of cells undergoing apoptosis increased monotonically with Caspase3 accumulation until 12 hours, following which even with an increase in *AUC_casp3_*, the rise in apoptosis was not substantial (50% to 80%) (Figure 4B(ii)). Similar behavior was observed for the TNFα+TPL case as well (Figure 4B(iii)). A comparison of the early phase apoptosis response for the TPL (Figure 4B(ii)) and TNFα+TPL (Figure 4B(iii)) cases shows marginally higher percentage apoptotic response as that in the former. This suggests that Caspase3 accumulation up to a particular threshold at the early time point (4-12h) is a pre-requisite to commit cells for apoptosis, which is only moderately affected by the Caspase3 surge at later time points (24h) for U937 cells. Next, we show the experimentally observed apoptosis levels with the model-predicted overall Caspase3 accumulation (Figure 4B, (iv)-(vi)). A comparison of Figure 4B(i)-(iii) and Figure 4B(iv)-(vi) demonstrates that the model simulated *AUC_casp3_* predicts the apoptosis levels as good as that by those from experimental measurements. This suggests that even though the accumulation information was not considered in the model training process (Figure 2B), the transients generated using the predicted model parameters adequately captures the apoptosis response via *AUC_casp3_*. In summary, both pJNK and pAKT, and their synergism with the NFκB regulation control the apoptotic response by modulating the overall Caspase3 levels.

### 5. Inhibiting pAKT and pJNK signaling alters Caspase3 transient response

In the previous section, we demonstrated that finetuning apoptotic response can be achieved by modulating Caspase3 transients. Since pAKT and pJNK transient correlates with the overall Caspase3 dynamics, inhibiting either of these marker proteins could essentially result in altered Caspase3 as well as apoptotic response. We effect these inhibitions by exposing the U937 cells to either Wortmannin (Wort), or SP600125 (SP6). Wort inhibits PI3K mediated pAKT activation (Teranishi et al., 2009; Zhang et al., 2009) as well as pJNK activation through MKK4/7 (Arcaro and Wymann, 1993; Ferby et al., 1994, 1996) (Figure 5A). On the other hand, SP6 affects both pJNK and pAKT activation via different pathways (Bennett et al., 2001; Moon et al., 2009; Mortenson et al., 2007) (Figure 5B). SP6 further activates Caspase3 (Moon et al., 2009; Yu et al., 2019) by down regulating Bcl2 activation and inhibits MAPK through the AP-1 and Fas driven pathway (Abreu-Martin et al., 1999; Bennett et al., 2001; Brandt et al., 2010; Eferl and Wagner, 2003; Mezosi et al., 2005; Shaulian and Karin, 2002). In order to investigate how the dynamic patterns of the entities altered in the presence of Wort or SP6 inhibitor, we phenomenologically incorporated their activity in our model, which will henceforth be referred to as *inhibitory model.* Details of inhibitory model are in Text S5, Supplementary Information.

**Figure 5.**
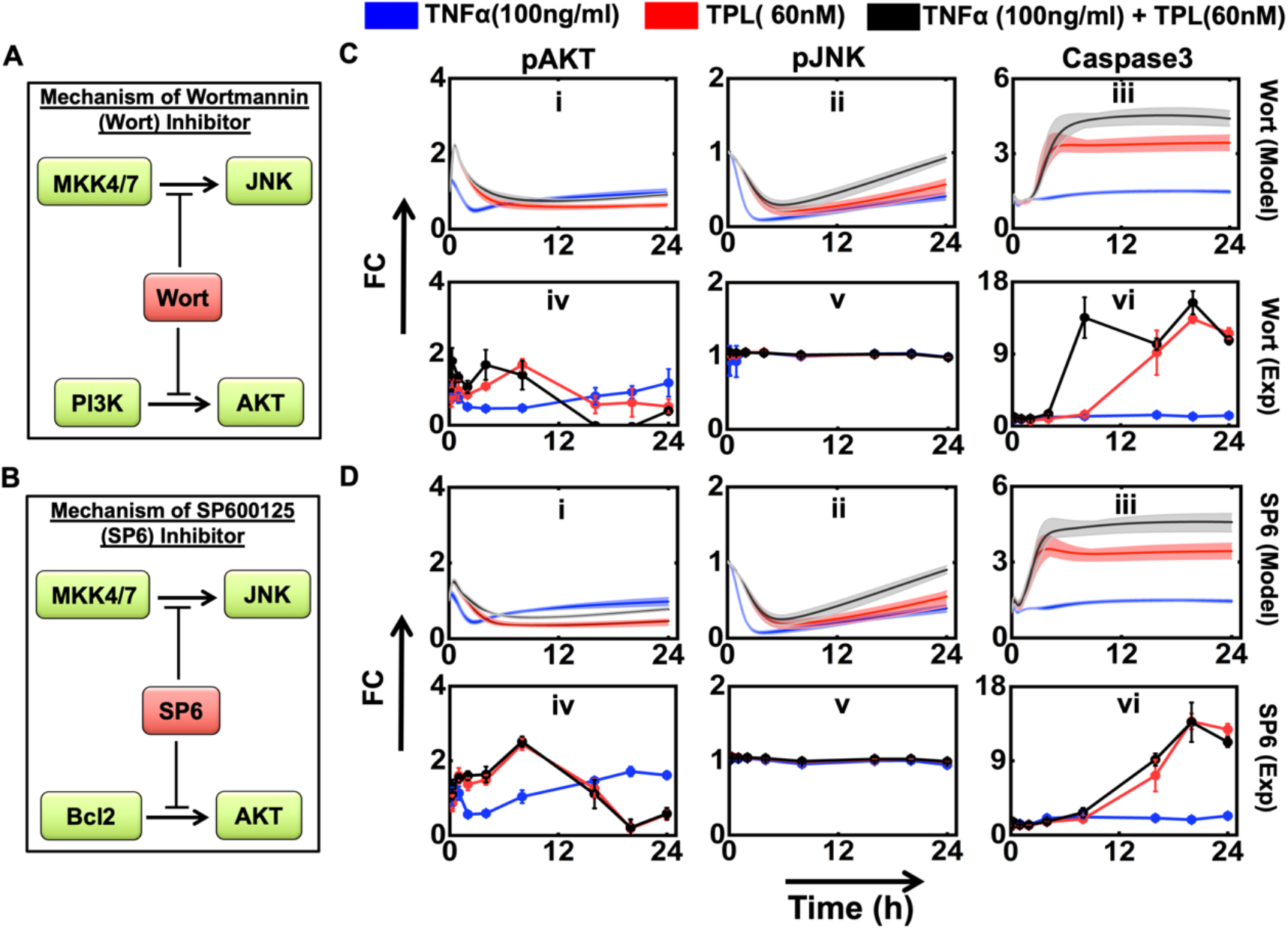
Model prediction and experimental validation of the marker proteins under the treatment of Wortmannin (Wort) and SP600125 (SP6) inhibitor. (A) and (B), respectively schematically represents the inhibitory action of Wort and SP6. (C) Effect of Wort inhibition on (i) pAKT, (ii) pJNK, and (iii) Caspase3 as predicted by model simulations. The corresponding experimental validations are in (iv), (v), and (vi), respectively. (D) Effect of SP6 inhibition on (i) pAKT, (ii) pJNK, and (iii) Caspase3 as predicted by model simulations. The corresponding experimental validations are in (iv), (v), and (vi), respectively. The rate constant values used for the inhibitory interactions are given in the caption of Figure S9, S10, Supplementary Information.

We first consider the case of Wort inhibition in our model simulations which demonstrate that pAKT levels decrease with time under all three stimulation conditions (Figure 5C(i)). This could be due to pJNK playing a significant role in activating pAKT (Figure 2A). On the contrary, for all three stimulation conditions, after an initial dip, pJNK dynamics showed a steady increasing trend (Figure 5C(ii)). For the case of only TNFα, in the presence of Wort, model predicts a marginal increase in Caspase3 level in the initial phase. However, when stimulated only with TPL or TNFα+TPL, simulations showed a steep increase in the Caspase3 levels in the initial phase. For all three stimulation conditions, the levels reached in the initial phase subsequently settles into the corresponding steady level till 24hr (Figure 5C(iii) and Figure S8). To validate these model predictions, we measured the transient levels of these marker proteins in Wort treated U937 cells under all three stimulation conditions. (Figure 5C(iv-vi)). (A comparison of the model predictions and the corresponding measurements with those obtained for the case of no Wort inhibition is presented in Figure S8.) The experimentally measured pAKT levels corroborated well with the theoretical model predictions (Figure 5C(i),(iv)). Caspase3 initial rise and subsequent saturation at a certain level predicted by the model for TPL and TNFα+TPL stimulation cases were qualitatively observed in experimentally detected dynamics (Figure 5C(iii), (iv)). The low Caspase3 levels predicted for TNFα treatment was also in line with that measured experimentally (Figure 5C(iii), (iv)). However, the sustained very low fold-change detected experimentally for pJNK could not be predicted by the model simulations (Figure 5C(ii), (v)).

We next investigate the effect of SP6 inhibition. The model predicted trend for all three marker proteins were similar to those obtained for the case of Wort inhibition for respective stimulation conditions (Figure 5D(i)-(iii)). (Inhibitory model predictions along with the experimental measurements under three different stimulation conditions are contrasted with those obtained for the case sans SP6 inhibition in Figure S8.) The experimentally measured dynamics of the marker proteins in SP6 treated U937 cells for all three stimulation conditions are in Figure 5D(iv)-(vi). These observations qualitatively substantiate the model predictions. Specifically, for the TPL and TNFα+TPL stimulation conditions, the rise in the Caspase3 levels were delayed as compared to model simulations (Figure 5D(iv), Figure 5D(iii)).

Flux analysis of the inhibitory model simulations (Text S5.3, Supplementary Information) revealed that in the presence of Wort, which arrests pAKT activation, pJNK primarily regulates the Caspase3 levels as it is relatively unchanged under all stimulation conditions (Figure S9, Supplementary Information). For the case of only TNFα stimulation, NFκB strongly inhibits the Caspase3 activation even in the absence of pAKT and pJNK due to Wort or SP6 treatment. Thus, the Caspase3 transient obtained is comparable for the cases of with (Figure 5C(iii), 5D(iii)) and without (Figure 2B(iii)) Wort/SP6 inhibition. However, under TPL and TNFα+TPL stimulation conditions, inhibition with Wort/SP6 results in an increased level of Caspase3 transients (Figure 5C(iii), 5D(iii)). This is because of the absence of the NFκB signaling mediated inhibitory action on Caspase3 as well due to the presence of TPL. Thus, the flux analysis identifies pJNK as the master regulator for Caspase3 in U937 cells.

Overall, our model could qualitatively predict the modulation in Caspase3 dynamic response under all the three experimental conditions (TNFα, TPL and TNFα+TPL) in presence of either Wort or SP6 inhibitor as substantiated by the experimental data.

### 6. Model adequately predicts apoptotic response in U937 cells under all stimulation conditions

We earlier demonstrated that the correlation of the model Caspase3 transients with apoptotic response quantitatively substantiated the corresponding experimental observations (Figure 4B). This suggests that the model is able to predict the cell-fate decision in U937 cells for which the *AUC_casp3_* can act as a suitable intracellular signature. Thus, fine-tuning apoptosis can be achieved by controlling *AUC_casp3_.* In order to test this hypothesis, we next contrast the model predicted apoptotic response with those observed experimentally when U937 cells are treated with Wort or SP6 inhibitors under all three stimulation conditions (TNFα, TPL, TNFα+TPL).

For predicting apoptosis level using the model simulated *AUC_casp3_* at a certain time, we assume the correlation between *AUC_casp3_* and apoptosis in Figure 4B(iv)-(vi) as a calibration map for the three stimulation conditions. In order to find the apoptotic response for the entire range of *AUC_casp3_* in Figure 4B(iv)-(vi), we quantify the correlation between model generated *AUC_casp3_* vs experimentally observed apoptosis using a polynomial curve fit. Details of the curve fitting are in Text S6, Supplementary information. Note that the curve fitting was performed using the 〈*AUC_casp3_*〉. Next, using the Wort or SP6 inhibitory model simulations, for a certain time duration under a specific stimulation condition, we estimated the 〈*AUC_casp3_*〉 (Methods), which was incorporated in the polynomial curve to arrive at the model predicted apoptosis percentage (Figure 6). The corresponding apoptosis percentage in U937 cells was measured experimentally (Methods).

**Figure 6:**
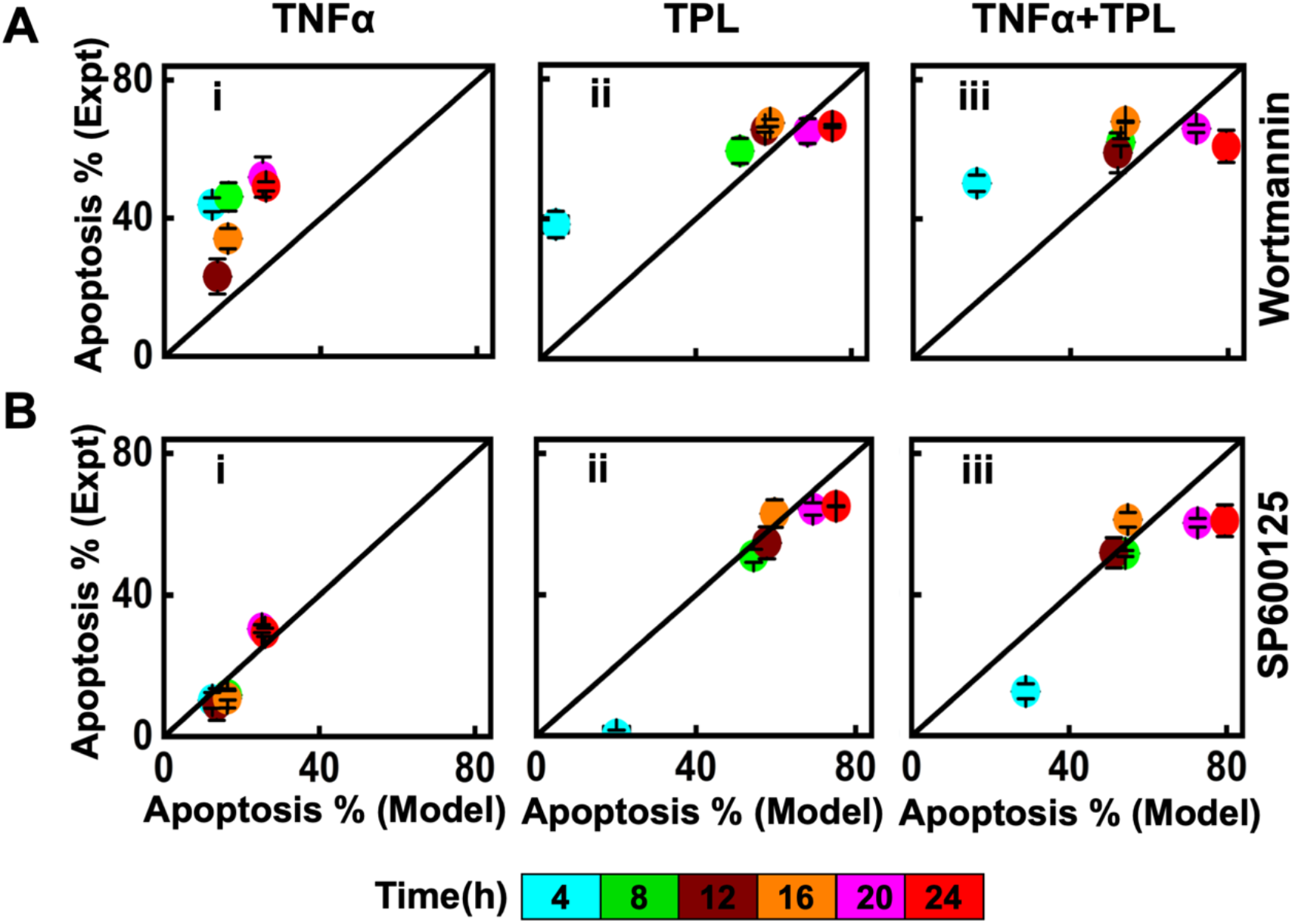
Comparison of experimental and model predicted apoptosis percentage under (A) Wort and (B) SP6 inhibitory conditions. (i), (ii), and (iii), respectively correspond to the predictions for TNFα, TPL, and TNFα+TPL stimulation conditions. While 5 best fit parameters were considered for model predictions, those of experiments are based on triplicates. The time points are appropriately color-coded in (B).

For the Wort inhibition case under only TNFα stimulation condition, the model underpredicted the experimentally observed apoptosis percentage (Figure 6A(i)). This is because of the Wort inhibitory model underestimating the pJNK level (Figure 5A(ii), blue). This deviation is reflected in the lower levels of Caspase3 transient (Figure 5A(iii), blue) causing decreased 〈*AUC_casp3_*〉. For the TPL and TNFα+TPL cases (Figure 6A(ii-iii)), the Wort inhibitory model predicts the experimental apoptosis percentage at all time durations other that 4 h. Note that after 16 h the experimental apoptosis levels become insensitive to the marginal increment in 〈*AUC_casp3_*〉 under both these conditions (Figure 5A(iii), red and black).

The SP6 inhibitory model predicted apoptosis levels match the experimental apoptotic measurements in U937 cells for all the time points during only TNFα stimulation (Figure 6B(i)). For the case of TPL and TNFα+TPL stimulations, for intermediate time points (8, 12, 16 h), the experimental observations corroborate model predictions (Figure 6B(ii),(iii)). However, for these two conditions, the model overpredicts the apoptosis levels at earlier and later duration. The over-prediction at the early time point (4 h) could be due to model predicting significantly higher levels of initial Caspase3 transients relative to the experimental observations (Figure 5B(iii), (iv)). Experimentally observed apoptosis response being insensitive to accumulated Caspase3 levels could explain the overprediction by the model at later time points (Figure 6B(ii),(iii), Figure 5B(vi), red and black).

Juxtaposition of model predicted 〈*AUC_casp3_*〉 and apoptosis leading to a correlation can help contrasting the experimentally observed and model simulation predicted apoptotic cell-fate response under different physiological conditions such as inhibition of an entity. Such a comparison can be achieved by inhibiting other entities in the network as well.

## Discussion

Dynamic crosstalk among intracellular signaling entities plays a crucial role in regulation of cell-fate as a response to the pleiotropic cytokine TNFα stimulation. In this study, we present an experimental data constrained mathematical model that can predict the apoptosis response under different physiological conditions related to TNFα signaling network. Using systems biology based approaches, we unravel the interdependence of pAKT and pJNK transient response and their synergistic modulation of apoptosis via Caspase3 dynamics in U937 cells. We demonstrated a systematic way to map the intracellular signaling response to the apoptotic cell-fate in a semi-quantitative manner.

TNFα signaling maintains a delicate balance between the survival and apoptotic response (Aggarwal et al., 2012). Our experimental findings unfolds that arresting survival signaling via NFκB pathway using Triptolide (TPL) leads to enhancement in apoptotic response in U937 cells (Figure 1). To distill out the intracellular signatures that may be controlling this response, we trained a 17 entity, 25 interaction TNFα network kinetic model using our high-throughput experimental transient data under TNFα, TPL, and TNFα+TPL stimulation conditions (Figure 2) for estimating the dynamic state and parameters. The model accurately predicted the experimentally observed transients for all three stimulation conditions.

The causal mechanism underlying the dynamic crosstalk governing the model trajectories was distilled out using the reaction flux analysis (Figure 3). We identified that pJNK transient response and PI3K play crucial roles in controlling pAKT dynamics. While PI3K primarily influences the late-phase activation of pAKT, pJNK affects it over the entire 24 h duration. However, pJNK transient is mostly affected by MKK4/7 and also inhibited by NFκB controlled XIAP/Gadd45B levels. Our flux analysis revealed that Caspase3 transients are regulated synergistically by both pAKT and pJNK levels in a stimulation dependent manner (Figure 3). When U937 is stimulated with TNFα, pJNK contribution to accumulated Caspase3 levels dominates over that by pAKT. However, the absence of survival signal due to the action of TPL establishes a balanced contribution by pJNK and pAKT towards accumulated Caspase3 levels (Figure 4). This shows that absence of NFκB activity improves the synergistic regulation of Caspase3 by pJNK and pAKT and thereby, enhance the apoptotic response (Figure 4). Our signal inhibition studies (Figure 5) further establish this synergism.

Correlation between the model simulated accumulated Caspase3 levels and experimentally detected apoptosis response enabled prediction of apoptotic phenotype in the presence of Wortmannin and SP600125 inhibitors for all three stimulation conditions (Figure 6). Even though the phenotypic predictions are valid only in the range of the accumulated Caspase3 levels, the systematic semi-quantitative approach employed can be easily extended to other perturbative conditions, which can be of ardent use for gaining insights into various therapeutic responses. Overall, we demonstrate that predicting the qualitative apoptotic phenotypic response is possible even without capturing the detailed pathway of apoptosis in an explicit manner in the model. However, there is scope for further improvement of the model to predict the experimental observations comprehensively.

## Conclusions

Using a systems biology experimental data driven mathematical model analysis, we unraveled the cross-talk between pJNK and pAKT transients in governing the dynamic apoptotic response via Caspase3. The synergism between the entities present in the TNFα network can be modulated by altering the physiological conditions such as inhibiting other pathway components. The model of the network can be extrapolated to mimic these conditions as well. However, apoptotic phenotype prediction using such an extended model would demand arriving at the associated correlation between accumulated Caspase3 and apoptosis levels, as this empirical relationship is context-specific. The model can be extended and improved upon in the future by incorporating stochasticity for predicting the cross-talk and phenotypic response at the single-cell level. We believe that the insights from the study can be capitalized upon to develop better therapeutic strategies involving the cytokine TNFα to shift the balance between survival, proliferation and apoptosis.

## Materials and Methods

### M1. Cell Culture and Reagents

U937 cells were procured from the Cell Repository at National Centre for Cell Science (NCCS), Pune, India. It was cultured in RPMI-1640, supplemented with 10% fetal bovine serum (FBS), 2 mmol/l L-glutamine, and 1% antibiotic–antimycotic solution, all procured from HiMedia (Mumbai, India). Cells were then collected, pelleted by centrifugation at 1,000 rpm for 5 min at room temperature (RT) and maintained at 37°C in a humidified 5% CO_2_ incubator. TNFα (Peprotech) was reconstituted in double-distilled water to a concentration of 100μg/ml. Triptolide (TPL) (Sigma) was dissolved in DMSO to a concentration of 1 mg/ml, the stock was stored at −20°C until use. Antibodies against different proteins ((phospho-pAKT(pS473)- Alexa 488-tagged antibody, phospho-pJNK(pT183/pY185)- PE tagged, and anti-active Caspase3- V450 tagged antibody)) were procured from BD Biosciences.

### M2. Apoptosis detection using Annexin V/PI staining

U937 cells were plated in 24-well plates. For a certain experimental condition, cells in different wells were treated with the corresponding stimulation for 4, 8, 12, 16, 20 or 24 h. For those conditions involving TPL, cells were pre-treated with it for 1 h prior to TNFα addition. After the completion of the stimulation time, cells were harvested, washed once with PBS, resuspended in 1X Annexin binding buffer and stained with FITC-labeled Annexin V and PI (BD Pharmingen, San Diego, CA, US). Cells were then analyzed on BD FACS Aria™ (BD Biosciences, San Jose, CA, US) within 30 min of dye addition. For every experimental condition and time point, three replicates were used.

### M3. Intracellular staining with Flowcytometry using Fluorescent Cell Barcoding (FCB)

U937 cells were seeded in 12 well plates for measuring signaling levels at 12 timepoints over a span of 24 h. For both TPL and TNFα+TPL stimulation conditions, cells were pre-treated with TPL for 1hr. After the stimulation, cells were fixed with 4% PFA for 10 min at RT, washed once with PBS and then transferred to a 96-well plate for further processing. Cells were then permeabilized with 500μl of 80% chilled methanol on ice for 1 h, and washed in PBS to remove leftover methanol. Now 50μl of 4 different concentrations (10μg/ml, 2μg/ml, 0.4μg/ml and 0.08μg/ml) of barcoding dye (Alexa-647) was added to the samples (450μl in PBS) making up the total volume to 500μl and kept on ice for 40min. After the incubation, cells were washed with 0.5%BSA in PBS to remove unbound dye. Finally, a combo tube was prepared that consisted of 4 different concentrations of the dye and washed thrice with 0.5%BSA in PBS to remove excess unbound dye. Cells resuspended in 0.5% BSA in PBS, were stained with antibodies against pAKT, pJNK and Caspase3 with dark incubation at RT for 30min. After removing the unbound antibody, cells resuspended in 300 μl of 0.5% BSA were analyzed for detecting fluorescence in various channels on a BD FACS Aria™ flow cytometer. A schematic of this entire high-throughput protocol is in Figure M1.

**Figure M1:**
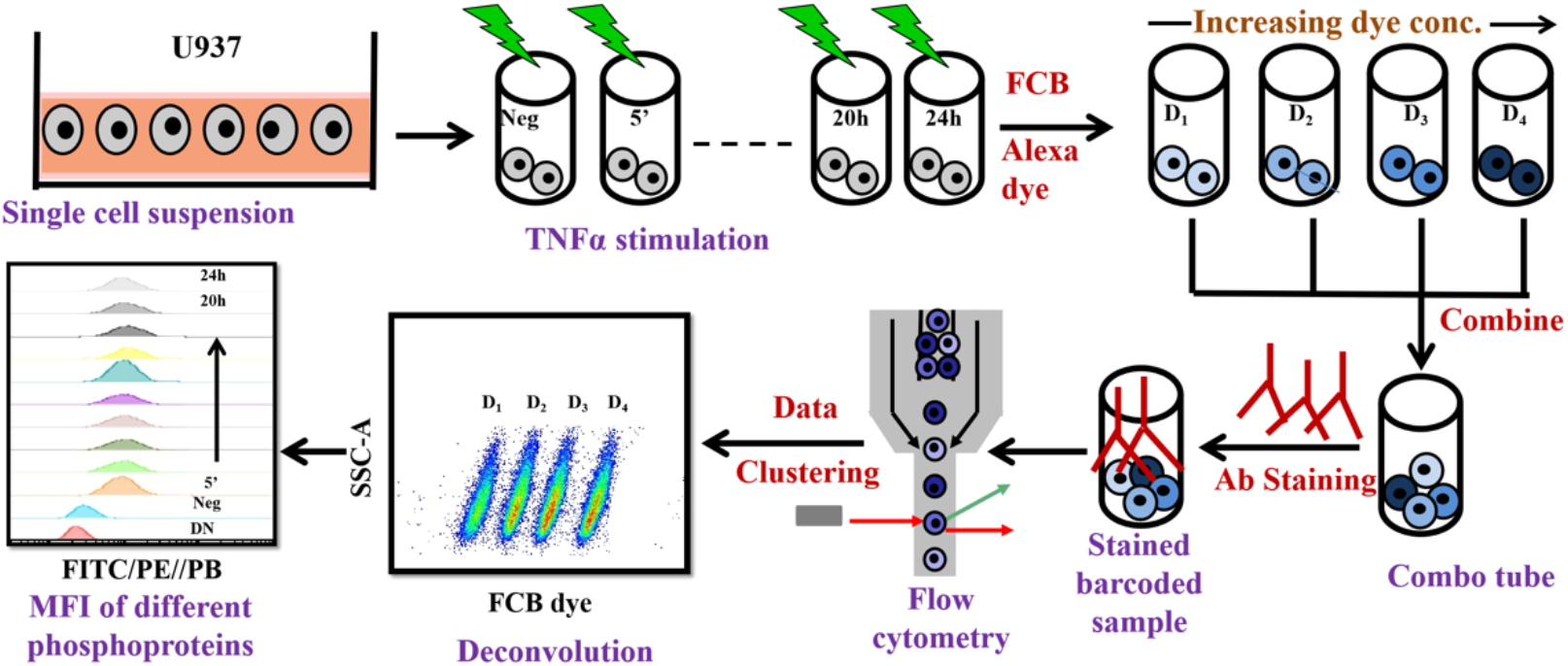
Schematic of high throughput signaling experimentation. Cells are harvested and incubated with inhibitor or stimulus for the desired time. After incubation, cells are fixed using paraformaldehyde and subsequently permeabilized with methanol. Four samples, each containing sufficient number of cells, are then incubated with amine-reactive fluorescent barcoding (FCB) (Krutzik and Nolan, 2006; Manohar et al., 2019) dye with different concentrations. These four samples are combined in a combo tube. After covalent binding, cells in the combo tube are washed several times to remove unbound dye, following which they are exposed to monoclonal antibody against the intracellular proteins. Fluorescence emitted by cells in the combo tube is then acquired on a flow cytometer. After acquisition, the samples are analyzed by gating and identifying individual samples displaying discrete fluorescent intensities (D1, D2, … D4) in the FCB channel. Mean fluorescent intensities of the fluorescence distribution corresponding to individual proteins are captured in different channels based on the fluorochrome antibody for further analysis.

### M4. Data quantification

The obtained file was then deconvoluted using FlowJo^®^ (version 10). Each fluorochrome used corresponds to a particular protein. Mean Fluorescence Intensity (MFI) of the distribution was used to calculate the relative fold change of the protein level upon stimulation. Relative fold change (FC) at a certain time point is given by

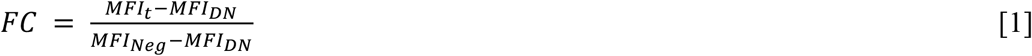

where, *MFI_t_*, *MFI_Neg_*, *MFI_DN_*, respectively represent MFI at a certain time *t*, negative control (no stimulation (0 h)), and double negative (only cells).

### M5. Signaling pathway perturbation with inhibitors

For each of the stimulation conditions, cells were treated with inhibitor Wortmannin (1μM) or SP600125 (10 μM) for 1 h prior to stimulation. After treatment, cells were stained as per the method specified earlier. Further the fluorescence emitted by the stained samples were acquired using BD FACS Aria™ following which FC at the measured time points were estimated using Eq. (1).

### M6. Model kinetic parameter estimation

The model kinetic parameter estimation was performed using the PottersWheel software (version 4.1.1) (Maiwald and Timmer, 2008) by employing “Trustregion” method with underlying ODEs integrated using CVODE method. The model system has been optimized ~2000 times by taking the same initial starting values of the parameters (parameters are globally fitted by considering a logarithmic parameter space) to fit the experimental data adequately. We considered χ^2^ value for a certain parameter set *P* as the fitting criteria, where χ^2^ is defined as

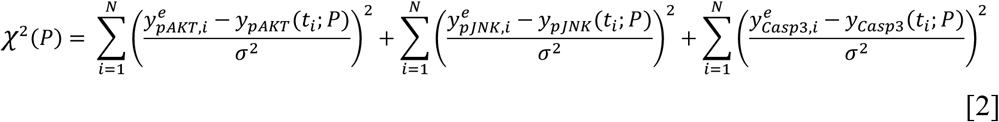

where, 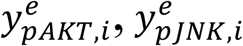 and 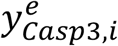 represent the mean of the triplicate experimental FC data for the pAKT, pJNK, Caspase3, respectively at the *i*th time point. *y_pAKT_*(*t_i_; P*), *y_pJNK_*(*t_i_; P*) and *y_Casp3_*(*t_i_; P*) are the levels of the respective proteins at the *i*th time point obtained from model simulations for a chosen parameter set *P*. *σ* is the standard deviation of the experimental data set. The detailed description of the model development is in Text S2, Supplementary Information.

### M7. Reaction Flux analysis

For a certain node such as pAKT, the rate law corresponding to every incoming and outgoing interactions were evaluated at every time point. For a certain interaction, a locus of these over 24 h is its reaction flux trajectory. We repeated this procedure to find the reaction flux trajectories for the interactions in which pAKT, pJNK and Caspase3 are involved. Details of these are in Text S4, Supplementary Information.

### M8. Correlation analysis and Linear regression

For a certain stimulation condition, area under the curve (*AUC*) was calculated for pAKT, pJNK and Caspase3 transients up to 8, 12, and 24 h time durations. For modelbased *AUC,* transients generated using 5 best parameter sets were employed. Linear regression (in R (Ritz and Streibig, 2005)) was carried out using the *AUC* from all 5 transients to find the constants *a* and *b* in overall pJNK and pAKT contributions in accumulated Caspase3 captured in

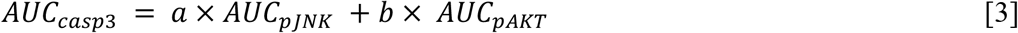

Relative contributions were estimated by normalizing all terms in Eq (3) with 〈*AUC_casp3_*〉 the average of *AUC_casp3_* across replicates. Thus, the average relative contributions of pAKT (*A_pAKT_*) and pJNK (*A_pJNK_*), respectively are defined as

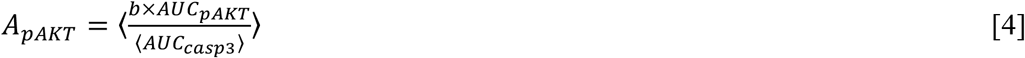

and

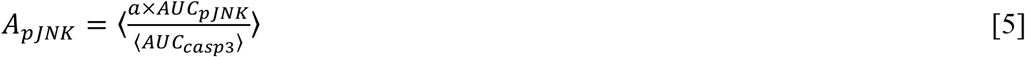

where, 〈. 〉 represents average across replicates. Note that *A_pJNK_* + *A_pAKT_* = 1.

## Acknowledgements

This study was funded by the grants CRG/2020/002672 (GV), MTR/2020/000589 (GV), CRG/2019/002640 (SK), and MTR/2020/000261 (SK) from Science and Engineering Research Board, Department of Science and Technology, Government of India. We gratefully acknowledge an access to the IIT Bombay FACS Central Facility and BSBE FACS Facility. SB and BT thank IIT Bombay for their fellowship.

## List of Supplementary Information

Text S1: Apoptosis and intracellular marker protein level detection

Text S2: Detailed model of the TNFα signaling network capturing the cross-talk between different entities

Text S3: Prediction of experimental dynamics by model simulations

Text S4: Quantifying the reaction fluxes influencing the dynamic levels of the three marker proteins.

Text S5: Model analysis under inhibitory conditions - Wortmannin (Wort) and SP600125 (SP6)

Text S6: Semi-quantitative relationship between 〈*AUC_casp3_*〉 and Apoptosis levels

## References

Abreu-Martin, M.T., Palladino, A.A., Faris, M., Carramanzana, N.M., Nel, A.E., and Targan, S.R. (1999). Fas activates the JNK pathway in human colonic epithelial cells: Lack of a direct role in apoptosis. Am. J. Physiol. - Gastrointest. Liver Physiol. 276, 599–605.

Adlung, L., Kar, S., Wagner, M.-C., She, B., Chakraborty, S., Bao, J., Lattermann, S., Boerries, M., Busch, H., Wuchter, P., et al. (2017). Protein abundance of AKT and ERK pathway components governs cell type-specific regulation of proliferation. Mol. Syst. Biol. 13, 904.

Aggarwal, B.B. (2003). Signalling pathways of the TNF superfamily: a double-edged sword. Nat. Rev. Immunol. 3, 745–756.

Aggarwal, B.B., Gupta, S.C., and Kim, J.H. (2012). Historical perspectives on tumor necrosis factor and its superfamily: 25 years later, a golden journey. Blood, J. Am. Soc. Hematol. 119, 651–665.

Aikin, R., Maysinger, D., and Rosenberg, L. (2004). Cross-talk between phosphatidylinositol 3-kinase/AKT and c-jun NH2-terminal kinase mediates survival of isolated human islets. Endocrinology 145, 4522–4531.

Amm, H.M., Oliver, P.G., Lee, C.H., Li, Y., and Buchsbaum, D.J. (2011). Combined modality therapy with TRAIL or agonistic death receptor antibodies. Cancer Biol. & Ther. 11, 431–449.

Van Antwerp, D.J., Martin, S.J., Kafri, T., Green, D.R., and Verma, I.M. (1996). Suppression of TNFα-induced apoptosis by NFκB. Science. 274, 787–789.

Arcaro, A., and Wymann, M.P. (1993). Wortmannin is a potent phosphatidylinositol 3-kinase inhibitor: the role of phosphatidylinositol 3, 4, 5-trisphosphate in neutrophil responses. Biochem. J. 296, 297–301.

Balkwill, F. (2009). Tumour necrosis factor and cancer. Nat. Rev. Cancer 9, 361–371.

Behar, M., and Hoffmann, A. (2010). Understanding the temporal codes of intracellular signals. Curr. Opin. Genet. & Dev. 20, 684–693.

Bennett, B.L., Sasaki, D.T., Murray, B.W., O’Leary, E.C., Sakata, S.T., Xu, W., Leisten, J.C., Motiwala, A., Pierce, S., Satoh, Y., et al. (2001). SP600125, an anthrapyrazolone inhibitor of Jun N-terminal kinase. Proc. Natl. Acad. Sci. 98, 13681–13686.

Beutler, B., Milsark, I.W., and Cerami, A.C. (1985). Passive immunization against cachectin/tumor necrosis factor protects mice from lethal effect of endotoxin. Science. 229, 869–871.

Bouwmeester, T., Bauch, A., Ruffner, H., Angrand, P.-O., Bergamini, G., Croughton, K., Cruciat, C., Eberhard, D., Gagneur, J., Ghidelli, S., et al. (2004). A physical and functional map of the human TNFα/NFκB signal transduction pathway. Nat. Cell Biol. 6, 97–105.

Brandt, B., Abou-Eladab, E.F., Tiedge, M., and Walzel, H. (2010). Role of the JNK/c-Jun/AP-1 signaling pathway in galectin-1-induced T-cell death. Cell Death & Dis. 1, e23–e23.

Burow, M.E., Weldon, C.B., Collins-Burow, B.M., Ramsey, N., McKee, A., Klippel, A., McLachlan, J.A., Clejan, S., and Beckman, B.S. (2000). Cross-talk between phosphatidylinositol 3-kinase and sphingomyelinase pathways as a mechanism for cell survival/death decisions. J. Biol. Chem. 275, 9628–9635.

Charles, K.A., Kulbe, H., Soper, R., Escorcio-Correia, M., Lawrence, T., Schultheis, A., Chakravarty, P., Thompson, R.G., Kollias, G., Smyth, J.F., et al. (2009). The tumor-promoting actions of TNFα involve TNFR1 and IL-17 in ovarian cancer in mice and humans. J. Clin. Invest. 119, 3011–3023.

Chen, G., and Goeddel, D. V (2002). TNF-R1 signaling: a beautiful pathway. Science. 296, 1634–1635.

Deng, Y., Ren, X., Yang, L., Lin, Y., and Wu, X. (2003). A JNK-dependent pathway is required for TNFα-induced apoptosis. Cell 115, 61–70.

Deppmann, C.D., and Janes, K.A. Cytokine-cytokine cross talk and cell-death decisions. Chap 8 in Systems Biology of Apoptosis, Editor Lavrik IN, (Springer), pp. 33–56.

Dhanasekaran, D.N., and Reddy, E.P. (2008). JNK signaling in apoptosis. Oncogene 27, 6245–6251.

Duffey, D.C., Crowl-Bancroft, C. V, Chen, Z., Ondrey, F.G., Nejad-Sattari, M., Dong, G., and Van Waes, C. (2000). Inhibition of transcription factor nuclear factor-κB by a mutant inhibitor-κBα attenuates resistance of human head and neck squamous cell carcinoma to TNFα caspase-mediated cell death. Br. J. Cancer 83, 1367–1374.

Eferl, R., and Wagner, E.F. (2003). AP-1: A double-edged sword in tumorigenesis. Nat. Rev. Cancer 3, 859–868.

Ferby, I.M., Waga, I., Sakanaka, C., Kume, K., and Shimizu, T. (1994). Wortmannin inhibits mitogen-activated protein kinase activation induced by platelet-activating factor in guinea pig neutrophils. J. Biol. Chem. 269, 30485–30488.

Ferby, I.M., Waga, I., Hoshino, M., Kume, K., and Shimizu, T. (1996). Wortmannin Inhibits Mitogen-activated Protein Kinase Activation by Platelet-activating Factor through a Mechanism Independent of p85/p110-type Phosphatidylinositol 3-Kinase. J. Biol. Chem. 271, 11684–11688.

Fouad, Y.A., and Aanei, C. (2017). Revisiting the hallmarks of cancer. Am. J. Cancer Res. 7, 1016–1036.

Galluzzi, L., Vitale, I., Aaronson, S.A., Abrams, J.M., Adam, D., Agostinis, P., Alnemri, E.S., Altucci, L., Amelio, I., Andrews, D.W., et al. (2018). Molecular mechanisms of cell death: recommendations of the Nomenclature Committee on Cell Death 2018. Cell Death & Differ. 25, 486–541.

Gerondakis, S., Grossmann, M., Nakamura, Y., Pohl, T., and Grumont, R. (1999). Genetic approaches in mice to understand Rel/NFκB and IκB function: transgenics and knockouts. Oncogene 18, 6888–6895.

Gierut, J.J., Wood, L.B., Lau, K.S., Lin, Y.-J., Genetti, C., Samatar, A.A., Lauffenburger, D.A., and Haigis, K.M. (2015). Network-level effects of kinase inhibitors modulate TNFα-induced apoptosis in the intestinal epithelium. Sci. Signal. 8, ra129.

Gregorc, V., De Braud, F.G., De Pas, T.M., Scalamogna, R., Citterio, G., Milani, A., Boselli, S., Catania, C., Donadoni, G., Rossoni, G., et al. (2011). Phase I study of NGR-hTNF, a selective vascular targeting agent, in combination with cisplatin in refractory solid tumors. Clin. Cancer Res. 17, 1964–1972.

Guo, Y.-L., Baysal, K., Kang, B., Yang, L.-J., and Williamson, J.R. (1998). Correlation between sustained c-Jun N-terminal protein kinase activation and apoptosis induced by tumor necrosis factor-α in rat mesangial cells. J. Biol. Chem. 273, 4027–4034.

Ha, S.-D., Martins, A., Khazaie, K., Han, J., Chan, B.M.C., and Kim, S.O. (2008). Cathepsin B is involved in the trafficking of TNFα containing vesicles to the plasma membrane in macrophages. J. Immunol. 181, 690–697.

Hanahan, D., and Weinberg, R.A. (2000). The hallmarks of cancer. Cell 100, 57–70.

Hanahan, D., and Weinberg, R.A. (2011). Hallmarks of cancer: the next generation. Cell 144, 646–674.

Herman, J.M., Wild, A.T., Wang, H., Tran, P.T., Chang, K.J., Taylor, G.E., Donehower, R.C., Pawlik, T.M., Ziegler, M.A., Cai, H., et al. (2013). Randomized phase III multi-institutional study of TNFerade biologic with fluorouracil and radiotherapy for locally advanced pancreatic cancer: final results. J. Clin. Oncol. 31, 886–894.

Hoffmann, A., and Baltimore, D. (2006). Circuitry of nuclear factor-κB signaling. Immunol. Rev. 210, 171–186.

Holoch, P.A., and Griffith, T.S. (2009). TNF-related apoptosis-inducing ligand (TRAIL): a new path to anti-cancer therapies. Eur. J. Pharmacol. 625, 63–72.

van Horssen, R., Ten Hagen, T.L.M., and Eggermont, A.M.M. (2006). TNFα in cancer treatment: molecular insights, antitumor effects, and clinical utility. Oncologist 11, 397–408.

Janes, K.A., Albeck, J.G., Gaudet, S., Sorger, P.K., Lauffenburger, D.A., and Yaffe, M.B. (2005). A systems model of signaling identifies a molecular basis set for cytokine-induced apoptosis. Science. 310, 1646–1653.

Jarosz-Griffiths, H.H., Holbrook, J., Lara-Reyna, S., and McDermott, M.F. (2019). TNF receptor signalling in autoinflammatory diseases. Int. Immunol. 31, 639–648.

Kandel, E.S., Skeen, J., Majewski, N., Di Cristofano, A., Pandolfi, P.P., Feliciano, C.S., Gartel, A., and Hay, N. (2002). Activation of Akt/protein kinase B overcomes a G2/M cell cycle checkpoint induced by DNA damage. Mol. Cell. Biol. 22, 7831–7841.

Kaufman, D.R., and Choi, Y. (1999). Signaling by tumor necrosis factor receptors: pathways, paradigms and targets for therapeutic modulation. Int. Rev. Immunol. 18, 405–427.

Kay, J., and Rahman, M.U. (2009). Golimumab: A novel human anti-TNFα monoclonal antibody for the treatment of rheumatoid arthritis, ankylosing spondylitis, and psoriatic arthritis. Core Evid. 4, 159–170.

Kearney, C.J., Vervoort, S.J., Hogg, S.J., Ramsbottom, K.M., Freeman, A.J., Lalaoui, N., Pijpers, L., Michie, J., Brown, K.K., Knight, D.A., et al. (2018). Tumor immune evasion arises through loss of TNF sensitivity. Sci. Immunol. 3, eaar3451.

Kist, M., and Vucic, D. (2021). Cell death pathways: intricate connections and disease implications. EMBO J. 40, e106700.

Kodama, S., Davis, M., and Faustman, D.L. (2005). The therapeutic potential of tumor necrosis factor for autoimmune disease: a mechanistically based hypothesis. Cell. Mol. Life Sci. C. 62, 1850–1862.

Kreuzaler, P., and Watson, C.J. (2012). Killing a cancer: what are the alternatives? Nat. Rev. Cancer 12, 411–424.

Krutzik, P.O., and Nolan, G.P. (2006). Fluorescent cell barcoding in flow cytometry allows high-throughput drug screening and signaling profiling. Nat. Methods 3, 361–368.

Lamb, J.A., Ventura, J.-J., Hess, P., Flavell, R.A., and Davis, R.J. (2003). JunD mediates survival signaling by the JNK signal transduction pathway. Mol. Cell 11, 1479–1489.

Lau, K.S., Cortez-Retamozo, V., Philips, S.R., Pittet, M.J., Lauffenburger, D.A., and Haigis, K.M. (2012). Multi-scale in vivo systems analysis reveals the influence of immune cells on TNFα-induced apoptosis in the intestinal epithelium. PLoS Biology 10, e1001393.

Leaner, V., Birrer, M.J., Rana, A., Shen, Y.H., Godlewski, J., Zhu, J., Sathyanarayana, P., and Tzivion, G. (2003). Cross-talk between JNK/SAPK and ERK/MAPK pathways: sustained activation of JNK blocks ERK activation by mitogenic factors. J. Biol. Chem. 278, 26715–26721.

Lee, K.Y., Chang, W., Qiu, D., Kao, P.N., and Rosen, G.D. (1999). PG490 (triptolide) cooperates with tumor necrosis factor-α to induce apoptosis in tumor cells. J. Biol. Chem. 274, 13451–13455.

Lipniacki, T., Puszynski, K., Paszek, P., Brasier, A.R., and Kimmel, M. (2007). Single TNFα trimers mediating NFκB activation: stochastic robustness of NFκB signaling. BMC Bioinformatics 8, 1–20.

Liu, T., Zhang, L., Joo, D., and Sun, S.-C. (2017). NFκB signaling in inflammation. Signal Transduct. Target. Ther. 2, 1–9.

Locksley, R.M., Killeen, N., and Lenardo, M.J. (2001). The TNF and TNF receptor superfamilies: integrating mammalian biology. Cell 104, 487–501.

Lopez, J., and Tait, S.W.G. (2015). Mitochondrial apoptosis: killing cancer using the enemy within. Br. J. Cancer 112, 957–962.

Lu, Y., Wang, X., Yan, W., Wang, H., Wang, M., Wu, D., Zhu, L., Luo, X., and Ning, Q. (2012). Liver TCRγδ+ CD3+ CD4- CD8- T cells contribute to murine hepatitis virus strain 3-induced hepatic injury through a TNFα dependent pathway. Mol. Immunol. 52, 229–236.

Maiwald, T., and Timmer, J. (2008). Dynamical modeling and multi-experiment fitting with PottersWheel. Bioinformatics 24, 2037–2043.

Manohar, S., Shah, P., Biswas, S., Mukadam, A., Joshi, M., and Viswanathan, G. (2019). Combining fluorescent cell barcoding and flow cytometry-based phospho-ERK1/2 detection at short time scales in adherent cells. Cytom. Part A 95, 192–200.

Mantovani, A., Allavena, P., Sica, A., and Balkwill, F. (2008). Cancer-related inflammation. Nature 454, 436–444.

Meffert, M.K., Chang, J.M., Wiltgen, B.J., Fanselow, M.S., and Baltimore, D. (2003). NFκB functions in synaptic signaling and behavior. Nat. Neurosci. 6, 1072–1078.

Mercogliano, M.F., Bruni, S., Mauro, F., Elizalde, P.V., and Schillaci, R. (2021). Harnessing Tumor Necrosis Factor Alpha to Achieve Effective Cancer Immunotherapy. Cancers (Basel). 13, 564.

Meyer, R., DAlessandro, L.A., Kar, S., Kramer, B., She, B., Kaschek, D., Hahn, B., Wrangborg, D., Karlsson, J., Kvarnstrom, M., et al. (2012). Heterogeneous kinetics of AKT signaling in individual cells are accounted for by variable protein concentration. Front. Physiol. 3, 451.

Mezosi, E., Wang, S.H., Utsugi, S., Bajnok, L., Bretz, J.D., Gauger, P.G., Thompson, N.W., and Baker, J.R. (2005). Induction and regulation of fas-mediated apoptosis in human thyroid epithelial cells. Mol. Endocrinol. 19, 804–811.

Micheau, O., and Tschopp, J. (2003). Induction of TNF receptor I-mediated apoptosis via two sequential signaling complexes. Cell 114, 181–190.

Moon, D.-O., Kim, M.-O., Kang, C.-H., Lee, J.-D., Choi, Y.H., and Kim, G.-Y. (2009). JNK inhibitor SP600125 promotes the formation of polymerized tubulin, leading to G 2/M phase arrest, endoreduplication, and delayed apoptosis. Exp. & Mol. Med. 41, 665–677.

Mortenson, M.M., Galante, J.G., Gilad, O., Schlieman, M.G., Virudachalam, S., Kung, H.-J., and Bold, R.J. (2007). BCL-2 functions as an activator of the AKT signaling pathway in pancreatic cancer. J. Cell. Biochem. 102, 1171–1179.

Newton, K., and Dixit, V.M. (2012). Signaling in innate immunity and inflammation. Cold Spring Harb. Perspect. Biol. 4, a006049.

Nurse, P. (2008). Life, logic and information. Nature 454, 424–426.

Oliver Metzig, M., Tang, Y., Mitchell, S., Taylor, B., Foreman, R., Wollman, R., and Hoffmann, A. (2020). An incoherent feedforward loop interprets NFκB/RelA dynamics to determine TNF-induced necroptosis decisions. Mol. Syst. Biol. 16, e9677.

Ozes, O.N., Akca, H., Gustin, J.A., Mayo, L.D., Pincheira, R., Korgaonkar, C.K., and Donner, D.B. (2005). Tumor Necrosis Factor-α/Receptor Signaling Through the Akt Kinase. In Cell Signaling in Vascular Inflammation, (Springer), pp. 13–22.

Pilati, P., Rossi, C.R., and Mocellin, S. (2008). Strategies to enhance the anticancer potential of TNF. Front Biosci 13, 3181–3193.

Purvis, J.E., and Lahav, G. (2013). Encoding and decoding cellular information through signaling dynamics. Cell 152, 945–956.

Rana, K., Liesveld, J.L., and King, M.R. (2009). Delivery of apoptotic signal to rolling cancer cells: A novel biomimetic technique using immobilized TRAIL and E-selectin. Biotechnol. Bioeng. 102, 1692–1702.

Reed, J.C., Miyashita, T., Takayama, S., Wang, H.-G., Sato, T., Krajewski, S., Aimé-Sempé, C., Bodrug, S., Kitada, S., and Hanada, M. (1996). Bcl-2 family proteins: regulators of cell death involved in the pathogenesis of cancer and resistance to therapy. J. Cell. Biochem. 60, 23–32.

Ritz, C., and Streibig, J.C. (2005). Bioassay analysis using R. J. Stat. Softw. 12, 1–22.

Sabio, G., and Davis, R.J. (2014). TNF and MAP kinase signalling pathways. In Seminars in Immunology, pp. 237–245.

Sankar, G., and Michael, K. (2002). Missing pieces in the NFκB puzzle. Cell 109, S81–96.

Sawada, M., Kiyono, T., Nakashima, S., Shinoda, J., Naganawa, T., Hara, S., Iwama, T., and Sakai, N. (2004). Molecular mechanisms of TNFα-induced ceramide formation in human glioma cells: p53-mediated oxidant stress-dependent and-independent pathways. Cell Death & Differ. 11, 997–1008.

Schleich, K., and Lavrik, I.N. (2013). Systems biology of death receptor-induced apoptosis. Chap2 in Systems Biology of Apoptosis, Editor Lavrik IN, (Springer), pp. 33–56.

Sharma, P., Wagner, K., Wolchok, J.D., and Allison, J.P. (2011). Novel cancer immunotherapy agents with survival benefit: recent successes and next steps. Nat. Rev. Cancer 11, 805–812.

Shaulian, E., and Karin, M. (2002). AP-1 as a regulator of cell life and death. Nat. Cell Biol. 4, E131–E136.

Sherekar, S., and Viswanathan, G.A. (2021). Boolean dynamic modeling of cancer signaling networks: Prognosis, progression, and therapeutics. Comput. Syst. Oncol. 1, e1017.

Shishodia, S., and Aggarwal, B.B. (2002). Nuclear factor-κB activation: a question of life or death. BMB Rep. 35, 28–40.

Son, M., Wang, A.G., Tu, H.-L., Metzig, M.O., Patel, P., Husain, K., Lin, J., Murugan, A., Hoffmann, A., and Tay, S. (2021). NFκB responds to absolute differences in cytokine concentrations. Sci. Signal. 14, eaaaz4382.

Steinman, L., Merrill, J.T., McInnes, I.B., and Peakman, M. (2012). Optimization of current and future therapy for autoimmune diseases. Nat. Med. 18, 59–65.

Takada, Y., and Aggarwal, B.B. (2004). TNF activates Syk protein tyrosine kinase leading to TNF-induced MAPK activation, NFκB activation, and apoptosis. J. Immunol. 173, 1066–1077.

Tang, D., Kang, R., Berghe, T. Vanden, Vandenabeele, P., and Kroemer, G. (2019). The molecular machinery of regulated cell death. Cell Res. 29, 347–364.

Teranishi, F., Takahashi, N., Gao, N., Akamo, Y., Takeyama, H., Manabe, T., and Okamoto, T. (2009). Phosphoinositide 3-kinase inhibitor (wortmannin) inhibits pancreatic cancer cell motility and migration induced by hyaluronan in vitro and peritoneal metastasis in vivo. Cancer Sci. 100, 770–777.

Ventura, J.-J., Hübner, A., Zhang, C., Flavell, R.A., Shokat, K.M., and Davis, R.J. (2006). Chemical genetic analysis of the time course of signal transduction by JNK. Mol. Cell 21, 701–710.

Vilcek, J. (2008). First demonstration of the role of TNF in the pathogenesis of disease. J. Immunol. 181, 5–6.

Vince, J.E., Wong, W.W.-L., Khan, N., Feltham, R., Chau, D., Ahmed, A.U., Benetatos, C.A., Chunduru, S.K., Condon, S.M., McKinlay, M., et al. (2007). IAP antagonists target cIAP1 to induce TNFα-dependent apoptosis. Cell 131, 682–693.

Vivanco, I., and Sawyers, C.L. (2002). The phosphatidylinositol 3-kinase-AKT pathway in human cancer. Nat. Rev. Cancer 2, 489–501.

Wajant, H., Pfizenmaier, K., and Scheurich, P. (2003). Tumor necrosis factor signaling. Cell Death & Differ. 10, 45–65.

Wallach, D., Varfolomeev, E.E., Malinin, N.L., Goltsev, Y. V, Kovalenko, A. V, and Boldin, M.P. (1999). Tumor necrosis factor receptor and Fas signaling mechanisms. Annu. Rev. Immunol. 17, 331–367.

Wang, L., Du, F., and Wang, X. (2008). TNFα induces two distinct caspase-8 activation pathways. Cell 133, 693–703.

Werner, S.L., Kearns, J.D., Zadorozhnaya, V., Lynch, C., O’Dea, E., Boldin, M.P., Ma, A., Baltimore, D., and Hoffmann, A. (2008). Encoding NFκB temporal control in response to TNF: distinct roles for the negative regulators IκBα and A20. Genes & Dev. 22, 2093–2101.

Wicovsky, A., Müller, N., Daryab, N., Marienfeld, R., Kneitz, C., Kavuri, S., Leverkus, M., Baumann, B., and Wajant, H. (2007). Sustained JNK activation in response to tumor necrosis factor is mediated by caspases in a cell type-specific manner. J. Biol. Chem. 282, 2174–2183.

Wu, G.S. (2009). TRAIL as a target in anti-cancer therapy. Cancer Lett. 285, 1–5.

Wullaert, A., Heyninck, K., and Beyaert, R. (2006). Mechanisms of crosstalk between TNF-induced NFκB and JNK activation in hepatocytes. Biochem. Pharmacol. 72, 1090–1101.

Yang, D., Tan, X., Lv, Z., Liu, B., Baiyun, R., Lu, J., and Zhang, Z. (2016). Regulation of Sirt1/Nrf2/TNFα signaling pathway by luteolin is critical to attenuate acute mercuric chloride exposure induced hepatotoxicity. Sci. Rep. 6, 1–12.

Yu, H., Wu, C.-L., Wang, X., Ban, Q., Quan, C., Liu, M., Dong, H., Li, J., Kim, G.-Y., Choi, Y.H., et al. (2019). SP600125 enhances C-2-induced cell death by the switch from autophagy to apoptosis in bladder cancer cells. J. Exp. & Clin. Cancer Res. 38, 1–13.

Zelová, H., and Hošek, J. (2013). TNFα signalling and inflammation: interactions between old acquaintances. Inflamm. Res. 62, 641–651.

Zhang, F., Zhang, T., Jiang, T., Zhang, R., Teng, Z., Li, C., Gu, Z.-P., and Mei, Q. (2009). Wortmannin potentiates roscovitine-induced growth inhibition in human solid tumor cells by repressing PI3K/Akt pathway. Cancer Lett. 286, 232–239.

Zhang, Q., Gupta, S., Schipper, D.L., Kowalczyk, G.J., Mancini, A.E., Faeder, J.R., and Lee, R.E.C. (2017). NFκB dynamics discriminate between TNF doses in single cells. Cell Syst. 5, 638–645.

Zhang, S., Lin, Z.-N., Yang, C.-F., Shi, X., Ong, C.-N., and Shen, H.-M. (2004). Suppressed NFκB and sustained JNK activation contribute to the sensitization effect of parthenolide to TNFα-induced apoptosis in human cancer cells. Carcinogenesis 25, 2191–2199.

