## Supplementary information for "Unraveling the intracellular cross-talk governing the balance between TNFα mediated survival and apoptosis signaling"

**Table of Contents**

**Text S1:** Apoptosis and intracellular marker protein level detection……………...[S4]
**Text S2:** Detailed model of the TNFα signaling network capturing the cross-talk between different entities……………………………………………………………..[S6]
**Text S3:** Prediction of experimental dynamics by model simulations…………...[S23]
**Text S4:** Quantifying the reaction fluxes influencing the dynamic levels of the three marker proteins…………………………………………………………………[S24]
**Text S5:** Model analysis under inhibitory conditions - Wortmannin (Wort) and SP600125 (SP6)……………………………………………………………………..[S26]
**Text S6:** Semi-quantitative relationship between $\langle AUC_{casp3} \rangle$ and Apoptosis…...[S31]
**References** ……………………………………………………………………[S32]

**List of Figures**

**Figure S1.** Fraction of U937 cell population stimulated with TNFα for 24 hrs undergoing apoptosis ……………………………………………..……………...[S4]
**Figure S2.** Time evolution of the histogram of fluorescence of pAKT, pJNK, and Caspase3 in U937 cells stimulated with TNFα………………………………………………………………….……………[S5]
**Figure S3**. Detailed TNFα signaling network……………………………………[S6]
**Figure S4**. Boxplot of the estimated kinetic parameters……………………….....[S21]
**Figure S5**. Model trajectories for best fitted parameter set with experimental measurements……………………………………………………………………..[S22]
**Figure S6.** Comparison of model predicted transients with experiment for TPL(10nM) in the presence and absence of TNFα (100ng/ml)…………………..[S23]
**Figure S7.** Flux analysis illustrates the dynamic influence of entities of TNFα network on the marker protein transients………………………………………[S25]
**Figure S8.** Comparison of the model and experimental dynamics of the marker proteins under (A) Wort or (B) SP6 inhibitory treatment for all three stimulation conditions. ………………………………………………………………………[S27]
**Figure S9.** Evolution of fluxes from important entities controlling the dynamics of pAKT, pJNK and Caspase3 under different experimental conditions in the presence and absence of Wort inhibitor …………………………………...…………[S28]
**Figure S10.** Evolution of fluxes from important entities controlling the dynamics of pAKT, pJNK and Caspase3 under different experimental conditions in the presence

and absence of SP6 inhibitor ………………………………………………………..[S29]

**List of Tables**

**Table S1:** Summary of the interactions in the TNFα signaling network along with the corresponding biochemical mode of action. Arrows and hammers, respectively represent activation and inhibition……………………..………………………..[S8]

**Table S2**: Ordinary differential equations of the TNFα signaling network and associated algebraic relations ……………………………………………………[S10]

**Table S3:** Description of the entities in the TNFα signaling network model and its state ……………………………………………………………………...……..[S13]

**Table S4:** Definition of the kinetic parameters involved in the model..................[S15]

**Table S5:** Reaction fluxes affecting the pJNK, pAKT and Caspase3 dynamics…[S24]

**Table S6:** Coefficients of the fourth order polynomial (Eq S6.1) under different stimulation conditions …………………………………………………………[S31]

### Text S1: Apoptosis and intracellular marker protein level detection

<u>S1.1: Apoptosis detection</u>

Apoptotic cells display phosphatidylserine (PS) on their membrane surface. Annexin-V conjugated with FITC has high binding affinity towards PS, thus cells which are programmed to undergo cell death will show FITC positive. However, cells with intact cellular membrane will repel the PI stain. Thus, the live quadrant shows both FITC and PE negative as cells are live and will not display PS on its cell surface membrane. Early apoptotic cells will display PS on its surface but have intact cellular membrane thus will show FITC positive and PE negative. Late apoptotic cells will display both PS on its surface as well will have disrupted cellular membrane showing both FITC and PE positive. Finally, necrotic cells will show no display of PS on its surface but have disrupted cellular membrane hence will only show PE positive. In Figure S1, we show a sample four-quadrant Annexin-V vs PI plot corresponding to the population of U937 cells, stimulated with TNFα for 24 hr, distributed among these quadrants. We considered the sum of both Early and Late apoptotic cells (quadrants 2 and 3) as the total percentage of cells undergoing apoptosis. Compensation controls included were untreated-unstained-cells, untreated cells stained with both Annexin-V and PE, treated cells stained only with Annexin-V and treated cells stained only with PE.

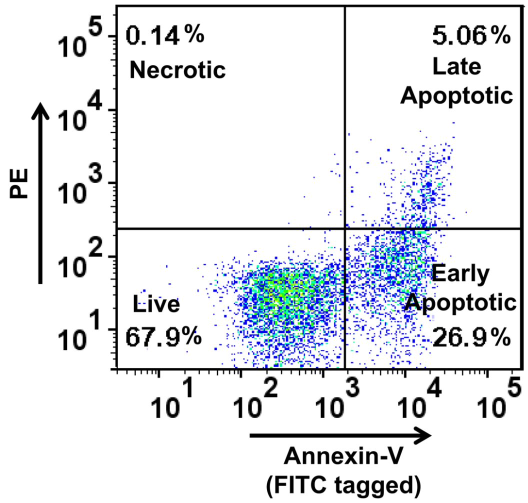


**Figure S1. Fraction of U937 cell population stimulated with TNFα for 24 hrs undergoing apoptosis.** The four quadrant plot depicting cells at different stages of cell death. Annexin-V is tagged with FITC and PI dye is detected in PE channel.

S1.2: Intracellular protein marker detection

The three marker proteins pAKT, pJNK, and Caspase3 are detected by tagging them with respective fluorescent dye tagged monoclonal antibody and the emitted fluorescence at single-cell level was detected in the appropriate channel in a flow cytometer (BD FACS Aria™). Figure S2 shows the histograms of the fluorescence emitted by these three markers at various time points for U937 cells stimulated with TNFα. The median fluorescence intensity ($MFI$) is estimated from these histograms and employed for arriving at the relative fold change (FC) given by

$$FC = \frac{MFI_t - MFI_{DN}}{MFI_{Neg} - MFI_{DN}} \quad [S1.1]$$

which is Eq 1, Methods M4, Main text and is used for further analysis. Note that antibodies tagged to different fluorochrome were selected based on the emission and excitation spectra such that it showed minimum spectral overlap among the samples. However, to correct the spectral overlap, if any, five compensation controls were used. These are untreated-unstained-cells, untreated cells stained with all monoclonal antibodies tagged with respective fluorochrome and one sample each containing treated cells with individual monoclonal antibody tagged with the corresponding fluorochrome.

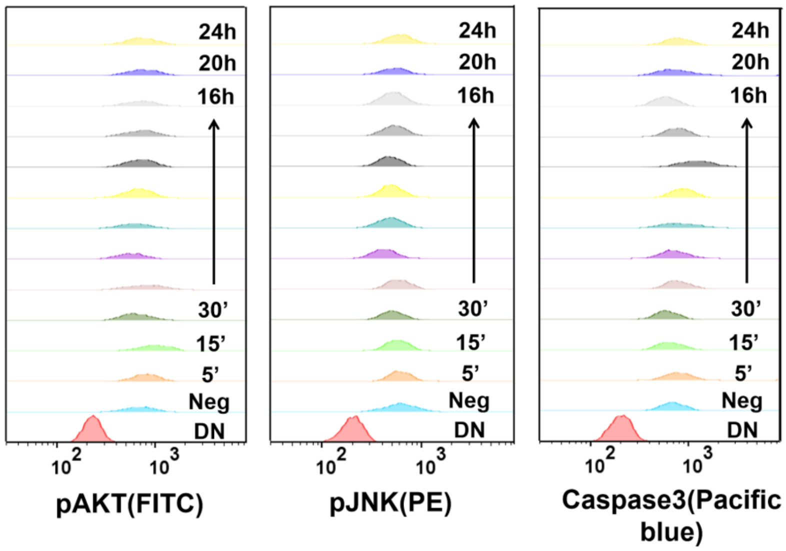


**Figure S2. Time evolution of the histogram of fluorescence of pAKT, pJNK, and Caspase3 in U937 cells stimulated with TNFα.** The emission from the fluorescence tagged monoclonal antibody against pAKT, pJNK and caspase3, respectively were detected in FITC, PE and Pacific blue channels.

### Text S2: Detailed model of the TNFα signaling network capturing the cross-talk between different entities

<u>S2.1: Detailed signaling network</u>

TNFα signaling network shown in Figure 2A consists of several interactions whose detailed underlying biochemical reaction along with the corresponding kinetic parameters are depicted in Figure S3 below. The biochemical details along with literature support is provided below. Moreover, procedure followed to develop the proposed model of the network based on these biochemical details is discussed in detail in this section.

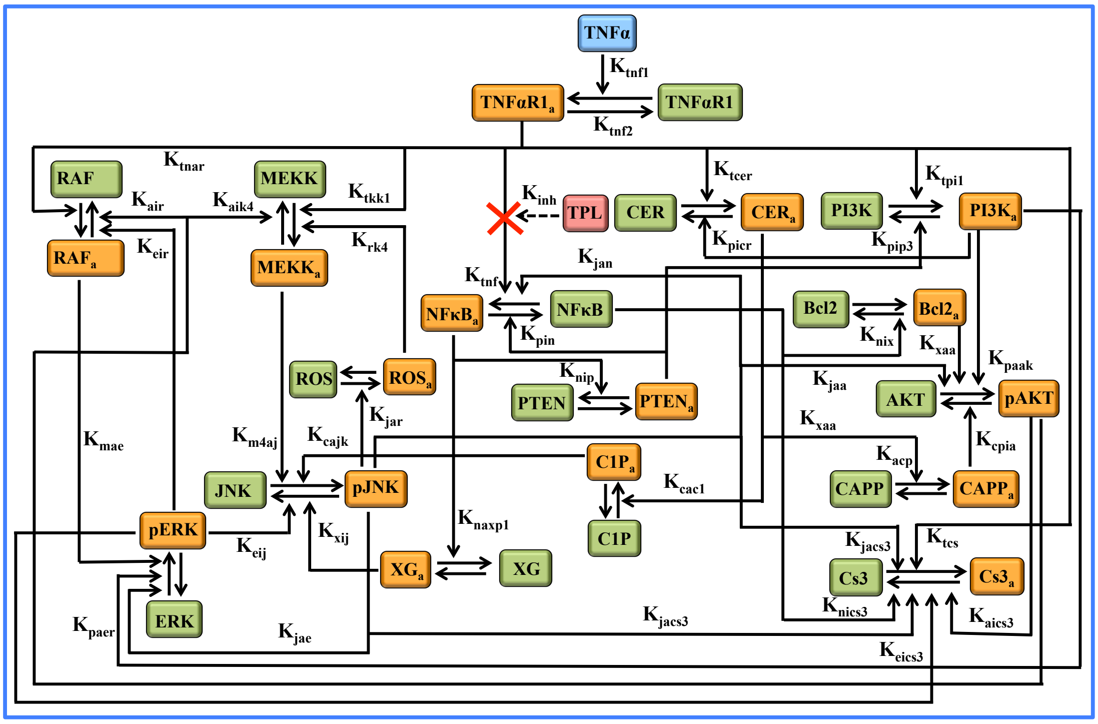


**Figure S3**. **Detailed TNFα signaling network**. While the green boxes represent the inactive form of an entity, their corresponding active form is denoted by orange box. The model contains total 33 species and total of 80 parameters including 3 scaling constants.

The interconnections have been selected based on the key players affecting the marker proteins (pAKT, pJNK, Caspase3) during TNFα stimulation. TNFα binds to its receptor TNFR1 transitioning it into its activated form $TNFR1_a$, which subsequently activates several downstream pathways. We included TNFα induced activation of RAF, MEKK, NFκB, Ceramide (Cer), PI3K, and Caspase3. The active forms RAFa and MEKKa further activate JNK to JNKa (Dhillon et al., 2007).

Further, RAFa also acts as a kinase to phosphorylate ERK to its active form pERK (Dhillon et al., 2007). Apart from MEKK assisted pJNK activation, active ceramide is involved in C1P mediated activation of pJNK (Gangoiti et al., 2008; Sawada et al., 2004). Similarly, apart from MEKK mediated pERK activation, PI3K too activates ERK (Grummisch et al., 2016). pERK negatively influences its own activation by inhibiting RAF activity (Dhillon et al., 2007). JNK, ROS and MEKK are locked in a positive feedback loop (Ventura et al., 2004). While pJNK could be inhibited by pERK (Monick et al., 2006), in turn, pERK is activated by pJNK. pERK is involved in inhibition of Caspase3 activation. This inhibition occurs via phosphorylation of Caspase9 at Thr 125 causing its inactivation (Allan et al., 2003). pJNK regulates Caspase3 activation which is modeled in a phenomenological manner as pJNK modulating apoptosis is context-dependent (Deng et al., 2003; Lamb et al., 2003).

In order to incorporate TNFα mediated cell-survival regulation, we considered pathways downstream of NFκB that includes both anti-apoptotic and survival signaling (Karin, 2002). As Triptolide (TPL) is an inhibitor of NFκB transactivation (Lee et al., 1999), we included pathways influenced by NFκB protein. The anti-apoptotic arm of NFκB signaling was phenomenologically introduced by its negative regulation of Caspase3 activity (Barkett and Gilmore, 1999). Apart from caspases, NFκB also influences pJNK and pAKT marker proteins. NFκB inhibits sustained activation of pJNK via XIAP-and-Gadd45B (XG) mediated inhibition (Papa et al., 2004). However, NFκB negatively and positively influences pAKT through different pathways. This activation is due to NFκB downregulating PTEN leading to increased pAKT levels via PI3K (Carracedo and Pandolfi, 2008; Vasudevan et al., 2004). On the other hand, NFκB transcriptionally inhibits Bcl2, which results in further inhibition of pAKT (Mortenson et al., 2007; Sohur et al., 1999). As PTEN is a well-known tumor suppressor gene, its action of inhibiting NFκB activity was included (Gustin et al., 2001). Active Ceramide negatively regulates pAKT via CAPP (Dobrowsky et al., 1993; Salinas et al., 2000). By phosphorylating pro-Caspase9 at Ser 196 leading to its inactivation, pAKT causes Caspase3 inhibition and subsequently results in negative regulation of apoptosis (Cardone et al., 1998). Moreover, pAKT downregulates the activated levels of RAF and MKK4 (Park et al., 2002; Rommel et al., 1999; Shaw and Kirshenbaum, 2006).

S2.2: Quantitative dynamic modeling of the TNFα signaling network

We model each of the interactions in Figure S3, listed in Table S1, with either law of mass action kinetics or phenomenological kinetics. Literature support for the mode of action chosen for the interactions are provided in Table S1. Using mass balance, we constructed the 16 variable ordinary differential equation based mathematical model of the TNFα network (Figure S3) constrained by a few algebraic conservation relations shown in Table S2. Those terms involving phenomenological relationship are discussed in section S2.3 below. Entities involved in the network are defined in Table S3. The definition of the parameters involved in the model are provided in Table S4.

**Table S1: Summary of the interactions in the TNFα signaling network along with the corresponding biochemical mode of action.** Arrows and hammers, respectively represent activation and inhibition.

| Sr. No | Interactions | Mode of Action | Description |
|---|---|---|---|
| 1. | $TNF \rightarrow TNFR1_a$ | Binding reaction | Binding of TNFα ligand with its receptor-1 (Wajant et al., 2003) |
| 2. | $TNF \rightarrow RAF$ | Enzymatic reaction | Raf complex gets activated via Phosphorylation at S338 and Y341. (Dhillon et al., 2007; Mason et al., 1999) |
| 3. | $TNF \rightarrow MEKK$ | Enzymatic reaction | MEKK1 leads to phosphorylated activation of MKK4 and MKK7 (Dhillon et al., 2007) |
| 4. | $TNF \rightarrow NF\kappa B$ | Enzymatic reaction | Activation of dimeric trancription factors (Karin, 2002) |
| 5. | $TNF \rightarrow CER$ | Catalysis reaction | Ceramide formation via activation of sphingomyelinases (Sawada et al., 2004) |
| 6. | $TNF \rightarrow PI3K$ | Enzymatic reaction | Activation of PI3K by TNFα (Ozes et al., 2005) |

| | | | |
|---|---|---|---|
| 7. | $RAF \rightarrow ERK$ | Enzymatic reaction | RAF-MEK-ERK is a RAS activated protein kinase cascade (Dhillon et al., 2007). |
| 8. | $ERK$ ⊣ RAF | Enzymatic reaction | ERK induced phosphorylation of B-RAF on T753 promotes the disassembly of RAF heterodimer. (Rushworth et al., 2006) |
| 9. | $MEKK \rightarrow JNK$ | Enzymatic reaction | MKK4 and MKK7 activates JNK by phosphorylating on Thr183 and Tyr185 residue (Dhillon et al., 2007). |
| 10. | $JNK \rightarrow ROS$ | Production reaction | TNFα induced pJNK mediated ROS production (Ventura et al., 2004). |
| 11. | $ROS \rightarrow MEKK$ | Enzymatic reaction | Increased level of ROS activates ASK1 which in turn activates MKK4 and MKK7 (Soga et al., 2012). |
| 12. | $JNK \rightarrow Caspase3$ | Catalytic and enzymatic reaction | JNK activates Caspase3 through a jBid-SMAC dependent mechanism (Deng et al., 2003). |
| 13. | $JNK \rightarrow AKT$ | Enzymatic reaction | Activation of AKT by pJNK through PDK-1 (Shaw and Kirshenbaum, 2006). |
| 14. | $JNK \rightarrow ERK$ | Enzymatic reaction | JNK indirectly activates $ERK$ through multiple interactive pathway |
| 15. | $ERK$ ⊣ JNK | Enzymatic reaction | pERK inhibits pJNK (Monick et al., 2006). |
| 16. | $NF\kappa B \rightarrow XG$ | Transcriptional activation | NFκB transcription factor induces transcription of anti-apoptotic proteins like XIAP and Gadd45B (Karin, 2002). |
| 17. | $JNK \rightarrow NF\kappa B$ | Enzymatic reaction | JNK indirectly activates $NF\kappa B$ through multiple interactive pathway |

| | | | |
|---|---|---|---|
| 18. | $XG \dashv$ JNK | Enzymatic reaction | pJNK activation is inhibited by NFκB regulated XIAP and Gadd45B (Papa et al., 2004). |
| 19. | $PTEN \dashv NF\kappa B$ | Enzymatic reaction | PTEN phosphatase inhibits NFκB activity (Gustin et al., 2001). |
| 20. | $NF\kappa B \dashv$ PTEN | Transcriptional reaction | NFκB inhibits PTEN (Vasudevan et al., 2004). |
| 21. | $PTEN \dashv$ PI3K | Enzymatic reaction | PTEN, a tumor supressor inhibits PI3K function (Carracedo and Pandolfi, 2008). |
| 22. | $NFkB \dashv$ Bcl2 | Transcriptional reaction | NFκB represses Bcl2 transcription (Sohur et al., 1999). |
| 23. | $NFkB \dashv$ Caspase3 | Multiple Inhibition | NFκB transcription factor induces transcription of various anti-apoptotic proteins (Barkett and Gilmore, 1999). |
| 24. | $ERK \dashv$ Caspase3 | Enzymatic reaction | pERK inhibits Caspase3 activation via phosphorylation of Caspase9 (Allan et al., 2003). |
| 25. | $AKT \dashv$ Caspase3 | Enzymatic reaction | pAKT inhibit cellular apoptosis by phosphorylating pro-Caspase9 at Ser 196 (Cardone et al., 1998). |

**Table S2: Ordinary differential equations of the TNFα signaling network and associated algebraic relations**

| *Ordinary differential equations* | |
|---|---|
| $\frac{dTNFR1_a}{dt} = \left((K_{base}) + (K_{tnf1} \times TNFR1 \times TNF) - (K_{tnf2} \times TNFR1_a)\right)$ | [1] |
| $\frac{dC1P_a}{dt} = ((K_{bc1} \times C1P) + (K_{cac1} \times C1P \times CER_a) - (K_{cac2} \times C1P_a))$ | [2] |
| $\frac{dXG_a}{dt} = \left((K_{bxg} \times XG) + (K_{naxp1} \times XG \times NFkB_a) - \left((K_{dxg} \times XG_a)\right)\right)$ | [3] |

| | |
|---|---|
| $\frac{dMEKK_a}{dt} = \left((K_{bkk1} \times MEKK) + (K_{tkk1} \times MEKK \times TNFR1_a) - (K_{dkk1} \times MEKK_a) + (K_{rak4} \times MEKK \times ROS_a) - (K_{aik4} \times MEKK_a \times pAKT)\right)$ | [4] |
| $\frac{dpJNK}{dt} = \left((K_{bjnk} \times JNK) + (K_{cajk} \times JNK \times C1P_a) - \left(\frac{K_{eij} \times pERK \times pJNK}{K_{eij1} + K_{eij2} \times pERK}\right) - (K_{djnk} \times pJNK) + (K_{m4aj} \times JNK \times MEKK_a) - (K_{xij} \times XG_a \times pJNK)\right)$ | [5] |
| $\frac{dNFkB_a}{dt} = \left(\left[\frac{(K_{bnf} \times NFkB)}{(1 + K_{inh} \times Tpl)}\right] + \left[\frac{(K_{tnf} \times NFkB \times TNFR1_a)}{(1 + K_{inh} \times Tpl)}\right] - (K_{pin} \times PTEN_a \times NFkB_a) + (K_{jan} \times pJNK \times NFkB) - (K_{dnf} \times NFkB_a)\right)$ | [6] |
| $\frac{dPI3K_a}{dt} = \left((K_{bp3k} \times PI3K) + (K_{tpi1} \times PI3K \times TNFR1_a) - (K_{pip3} \times PI3K_a \times PTEN_a) - (K_{bdp3k} \times PI3K_a)\right)$ | [7] |
| $\frac{dPTEN_a}{dt} = \left((K_{bpt} \times PTEN) - (K_{nip} \times NFkB_a \times PTEN_a) - (K_{dpt} \times PTEN_a)\right)$ | [8] |
| $\frac{dpAKT}{dt} = \left((K_{bak} \times AKT) + (K_{paak} \times PI3K_a \times AKT) - (K_{bdak} \times pAKT) + (K_{jaa} \times pJNK \times AKT) - (K_{cpia} \times CAPP_a \times pAKT) + (K_{xaa} \times Bcl2_a \times AKT)\right)$ | [9] |
| $\frac{dCER_a}{dt} = \left((K_{bcer} \times CER) + (K_{tcr} \times CER \times TNFR1_a) - (K_{dcer} \times CER_a) - (K_{picr} \times PI3Ka \times CER_a)\right)$ | [10] |
| $\frac{dCAPP_a}{dt} = \left((K_{bcpp} \times CAPP) + (K_{acp} \times CAPP \times CER_a) - (K_{dcpp} \times CAPP_a)\right)$ | [11] |

| | |
|---|---|
| $\frac{dBcl2_a}{dt} = \big((K_{bx} \times Bcl2) - (K_{nix} \times Bcl2_a \times NFkB_a) - (K_{dx} \times Bcl2_a)\big)$ | [12] |
| $\frac{dROS_a}{dt} = \big((K_{bros} \times ROS) + (K_{jar} \times ROS \times pJNK) - (K_{dros} \times ROS_a)\big)$ | [13] |
| $\frac{dRAF_a}{dt} = \big((K_{braf} \times RAF) + (K_{tnar} \times TNFR1_a \times RAF) - (K_{dbraf} \times RAF_a) - (K_{eir} \times RAF_a \times pERK) - (K_{air} \times RAF_a \times pAKT)\big)$ | [14] |
| $\frac{dpERK}{dt} = \big((K_{berk} \times ERK) + (K_{mae} \times ERK \times RAF_a) - (K_{dbrk} \times pERK) + (K_{paer} \times ERK \times PI3K_a) + (K_{jae} \times ERK \times pJNK)\big)$ | [15] |
| $\frac{dCs3_a}{dt} = \left(\left[\frac{(K_{jacs3} \times Cs3 \times pJNK)}{(K_{jac2} + (n1 \times pJNK))}\right] - \left[\frac{(K_{eics3} \times Cs3_a \times pERK)}{(K_{eic1} + (K_{eic2} \times pERK))}\right] - \left[\frac{(K_{aics3} \times Cs3_a \times pAKT^{K_{n2}})}{(K_{aic1} + (K_{aic2} \times pAKT^{K_{n2}}))}\right] + (K_{bcs3} \times Cs3) + (K_{tacs} \times Cs3 \times TNFR1_a) - (K_{nics3} \times Cs3_a \times NFkB_a) - (K_{dbcs3} \times Cs3_a)\right)$ | [16] |
| *Algebraic Relations* | |
| $TNFR1 = (TNFR1T - TNFR1_a)$ | [1] |
| $CER = (CERT - CER_a)$ | [2] |
| $XG = (XGT - XG_a)$ | [3] |
| $MEKK = (MEKKT - MEKK_a)$ | [4] |
| $JNK = (JNKT - pJNK)$ | [5] |
| $NFkB = (NFkBT - NFkB_a)$ | [6] |
| $PI3K = (PI3KT - PI3K_a)$ | [7] |
| $PTEN = (PTENT - PTEN_a)$ | [8] |

| | |
|---|---|
| $AKT = (AKTT - pAKT)$ | [9] |
| $CER = (CERT - CER_a)$ | [10] |
| $CAPP = (CAPPT - CAPP_a)$ | [11] |
| $Bcl2 = (Bcl2T - Bcl2_a)$ | [12] |
| $ROS = (ROST - ROS_a)$ | [13] |
| $RAF = (RAFT - RAF_a)$ | [14] |
| $ERK = (ERKT - pERK)$ | [15] |
| $Cs3 = (Cs3T - Cs3_a)$ | [16] |
| $baAKT = \left(\frac{K_{bak} \times AKTT}{K_{bak} + K_{bdak}}\right)$ | [17] |
| $baJNK = \left(\frac{K_{bjnk} \times JNKT}{K_{bjnk} + K_{bjnk}}\right)$ | [18] |
| $baCs3 = \left(\frac{K_{bcs3} \times Cs3T}{K_{bcs3} + K_{dbcs3}}\right)$ | [17] |
| *Observables* | |
| $FC\_AKT = S1 \times \left(\frac{pAKT}{baAKT}\right)$ | [1] |
| $FC\_JNK = S \times \left(\frac{pJNK}{baJNK}\right)$ | [2] |
| $FC\_Cs3 = S2 \times \left(\frac{Cs3_a}{baCs3}\right)$ | [3] |

**Table S3: Description of the entities in the TNFα signaling network model and its state**

| Symbol | Descriptions of the species | |
|---|---|---|
| $TNF$ | The TNF-α ligand | [1] |
| $TNFR1$ | The TNF-α receptor-1 | [2] |
| $TNFR1_a$ | Active complex formed by binding of TNF-α ligand with the TNF-α receptor-1 | [3] |
| $C1P$ | Inactive form of C1P protein | [4] |
| $C1P_a$ | Phosphorylated form of C1P protein | [5] |
| $XG$ | Inactive form of XIAP and Gadd45B protein | [6] |
| $XG_a$ | Active form of XIAP and Gadd45B protein | [7] |

| | | |
|---|---|---|
| $MEKK$ | Inactive form of MKK4 and MKK7 protein | [8] |
| $MEKK_a$ | Phosphorylated form of MKK4 and MKK7 protein | [9] |
| $JNK$ | Inactive form of JNK protein | [10] |
| $pJNK$ | Phosphorylated form of JNK protein | [11] |
| $NF\kappa B$ | Inactive form of NFκB protein | [12] |
| $NF\kappa B_a$ | Active form of NFκB protein | [13] |
| $Tpl$ | Triptolide, inhibitor of NFκB protein | [14] |
| $PTEN$ | Inactive form of PTEN protein | [15] |
| $PTEN_a$ | Active form of PTEN protein | [16] |
| $PI3K$ | Inactive form of PI3K kinase | [17] |
| $PI3K_a$ | Active form of PI3K kinase | [18] |
| $AKT$ | Inactive form of AKT protein | [19] |
| $pAKT$ | Phosphorylated form of AKT protein | [20] |
| $CER$ | Inactive form of CERAMIDE protein | [21] |
| $CER_a$ | Active form of CERAMIDE protein | [22] |
| $CAPP$ | Inactive form of CAPP protein | [23] |
| $CAPP_a$ | Active form of CAPP protein | [24] |
| $Bcl2$ | Inactive form of Bcl2 protein | [25] |
| $Bcl2_a$ | Active form of Bcl2 protein | [26] |
| $RAF$ | Inactive form of RAF complex | [27] |
| $RAF_a$ | Phosphorylated form of RAF complex | [28] |
| $ROS$ | Inactive form of ROS protein | [29] |
| $ROS_a$ | Active form of ROS protein | [30] |
| $ERK$ | Inactive form of ERK1/2 protein | [31] |
| $pERK$ | Phosphorylated form of ERK1/2 protein | [32] |
| $Cs3$ | Inactive form of cleaved Caspase-3 protein | [33] |
| $Cs3_a$ | Active form of cleaved Caspase-3protein | [33] |

**Table S4: Definition of the kinetic parameters involved in the model.**

| Symbol | Descriptions of the Parameter | Unit | |
|---|---|---|---|
| $K_{base}$ | Basal synthesis rate of the TNF-α receptor-1 | $h^{-1}$ | [1] |
| $K_{tnf1}$ | Complex ($TNFR1_a$) formation rate or Binding rate of TNF-α ligand with TNF-α receptor-1 | $nM^{-1}$ $h^{-1}$ | [2] |
| $K_{tnf2}$ | Dissociation rate of the active Complex ($TNFR1_a$) | $h^{-1}$ | [3] |
| $K_{bc1}$ | Basal phosphorylation rate of the C1P protein | $h^{-1}$ | [4] |
| $K_{cac1}$ | $CER_a$ mediated phosphorylation rate of C1P protein | $nM^{-1}$ $h^{-1}$ | [5] |
| $K_{cac2}$ | Dephosphorylation rate of C1P protein | $h^{-1}$ | [6] |
| $K_{bxg}$ | Basal activation rate of the XG protein | $h^{-1}$ | [7] |
| $K_{naxp1}$ | $NF\kappa B_a$ mediated transcriptional activation rate of XG protein | $nM^{-1}$ $h^{-1}$ | [8] |
| $K_{dxg}$ | Deactivation rate of XG protein | $h^{-1}$ | [9] |
| $K_{bkk1}$ | Basal phosphorylation rate of the MEKK protein | $h^{-1}$ | [10] |
| $K_{tkk1}$ | $TNFR1_a$ mediated phosphorylation rate of MEKK protein | $nM^{-1}$ $h^{-1}$ | [11] |
| $K_{dkk1}$ | Dephosphorylation rate of MEKK protein | $h^{-1}$ | [12] |
| $K_{rk4}$ | $ROS_a$ mediated phosphorylation rate of MEKK protein | $nM^{-1}$ $t^{-1}$ | [13] |
| $K_{aik4}$ | pAKT mediated dephosphorylation rate of MEKK protein | $nM^{-1}$ $h^{-1}$ | [14] |
| $K_{bjnk}$ | Basal phosphorylation rate of the JNK protein | $h^{-1}$ | [15] |
| $K_{cajk}$ | $C1P_a$ mediated phosphorylation of JNK protein | $nM^{-1}$ $h^{-1}$ | [16] |
| $K_{eij}$ | pERK mediated deactivation rate of JNK protein | $nM^{-1}$ $h^{-1}$ | [17] |
| $K_{eij1}$ | Constant controlling the pERK thresholding effect on pJNK inhibition | - | [18] |

| | | | |
|---|---|---|---|
| $K_{eij2}$ | Constant capturing the saturation effect of pERK on pJNK inhibition | $nM^{-1}$ | [19] |
| $K_{djnk}$ | Dephosphorylation rate of JNK protein | $h^{-1}$ | [20] |
| $K_{m4aj}$ | MEKK$_a$ mediated phosphorylation of JNK protein | $nM^{-1}$ $h^{-1}$ | [21] |
| $K_{xij}$ | XG$_a$ mediated deactivation rate of JNK protein | $nM^{-1}$ $h^{-1}$ | [22] |
| $K_{bnf}$ | Basal activation rate of the NFκB protein | $h^{-1}$ | [23] |
| $K_{inh}$ | Coefficient controlling the Tpl mediated NFκB inhibition | $nM^{-1}$ | [24] |
| $K_{bnf}$ | TNFR1$_a$ mediated activation rate of NFκB protein | $nM^{-1}$ $h^{-1}$ | [25] |
| $K_{jan}$ | pJNK mediated activation rate of NFκB protein | $nM^{-1}$ $h^{-1}$ | [26] |
| $K_{pin}$ | PTEN$_a$ mediated deactivation rate of NFκB protein | $nM^{-1}$ $h^{-1}$ | [27] |
| $K_{dnf}$ | Deactivation rate of NFκB protein | $h^{-1}$ | [28] |
| $K_{bpt}$ | Basal phosphorylation rate of the PTEN protein | $h^{-1}$ | [29] |
| $K_{nip}$ | NFκB$_a$ mediated inhibition rate of PTEN protein | $nM^{-1}$ $h^{-1}$ | [30] |
| $K_{dpt}$ | Deactivation rate of PTEN protein | $h^{-1}$ | [31] |
| $K_{bp3k}$ | Basal phosphorylation rate of PI3K protein | $h^{-1}$ | [32] |
| $K_{tpi1}$ | TNFR1$_a$ mediated activation rate of PI3Ka protein | $nM^{-1}$ $h^{-1}$ | [33] |
| $K_{pip3}$ | PTEN$_a$ mediated dephosphorylation rate of PI3K protein | $nM^{-1}$ $h^{-1}$ | [33] |
| $K_{bdp3k}$ | Dephosphorylation rate of PI3K protein | $h^{-1}$ | [34] |
| $K_{bak}$ | Basal phosphorylation rate of AKT protein | $h^{-1}$ | [35] |
| $K_{paak}$ | PI3K$_a$ mediated phosphorylation rate of AKT protein | $nM^{-1}$ $h^{-1}$ | [36] |

| | | | |
|---|---|---|---|
| $K_{jaa}$ | pJNK mediated activation rate of AKT protein | $nM^{-1}$ $h^{-1}$ | [37] |
| $K_{cpia}$ | $CAPP_a$ mediated dephosphorylation rate of AKT protein | $nM^{-1}$ $h^{-1}$ | [38] |
| $K_{xaa}$ | $Bcl2_a$ mediated phosphorylation rate of AKT protein | $nM^{-1}$ $h^{-1}$ | [39] |
| $K_{bdak}$ | Dephosphorylation rate of AKT protein | $h^{-1}$ | [40] |
| $K_{bcer}$ | Basal phosphorylation rate of CERAMIDE protein | $h^{-1}$ | [41] |
| $K_{tcr}$ | $TNFR1_a$ mediated catalysis rate of CERAMIDE protein | $nM^{-1}$ $h^{-1}$ | [42] |
| $K_{picr}$ | $PI3K_a$ mediated inhibition rate of CERAMIDE protein | $nM^{-1}$ $h^{-1}$ | [43] |
| $K_{dcer}$ | Deactivation rate of CERAMIDE protein | $h^{-1}$ | [44] |
| $K_{bx}$ | Basal activation rate of Bcl2 protein | $h^{-1}$ | [45] |
| $K_{dx}$ | Deactivation rate of Bcl2 protein | $h^{-1}$ | [46] |
| $K_{nix}$ | $NF\kappa B_a$ mediated inhibition rate of Bcl2 protein transcription | $nM^{-1}$ $h^{-1}$ | [47] |
| $K_{braf}$ | Basal phosphorylation rate of RAF protein | $h^{-1}$ | [48] |
| $K_{tnar}$ | $TNFR1_a$ mediate phosphorylation rate of RAF protein | $nM^{-1}$ $h^{-1}$ | [49] |
| $K_{eir}$ | pERK mediated phosphorylation rate of RAF protein leading to deactivation of RAF | $nM^{-1}$ $h^{-1}$ | [50] |
| $K_{air}$ | pAKT mediated phosphorylation rate of RAF protein leading to deactivation of RAF | $nM^{-1}$ $h^{-1}$ | [51] |
| $K_{dbraf}$ | Deactivation rate of RAF protein | $h^{-1}$ | [52] |
| $K_{bros}$ | Basal activation rate of ROS protein | $h^{-1}$ | [53] |
| $K_{jar}$ | pJNK mediated activation rate of ROS protein | $nM^{-1}$ $h^{-1}$ | [54] |
| $K_{dros}$ | Deactivation rate of ROS protein | $h^{-1}$ | [55] |
| $K_{berk}$ | Basal activation rate of ERK protein | $h^{-1}$ | [56] |

| | | | |
|---|---|---|---|
| $K_{mae}$ | $RAF_a$ mediated activation rate of ERK protein | $nM^{-1}$ $h^{-1}$ | [57] |
| $K_{paer}$ | $PI3K_a$ mediated activation rate of ERK protein | $nM^{-1}$ $h^{-1}$ | [58] |
| $K_{jae}$ | pJNK mediated activation rate of ERK protein | $nM^{-1}$ $h^{-1}$ | [59] |
| $K_{dbrk}$ | Deactivation rate of ERK protein | $h^{-1}$ | [60] |
| $K_{bcs3}$ | Basal activation rate of Cs3 protein | $h^{-1}$ | [61] |
| $K_{jacs3}$ | pJNK mediate activation rate of Cs3 protein | $h^{-1}$ | [62] |
| $K_{jac2}$ | Constant controlling the pJNK thresholding effect on Cs3 activation | - | [63] |
| $n1$ | Constant capturing the saturation effect of pJNK on Cs3 activation | $nM^{-1}$ | [64] |
| $K_{tcs}$ | $TNFR1_a$ mediate activation rate of Cs3 protein | $nM^{-1}$ $h^{-1}$ | [65] |
| $K_{nics3}$ | $NF\kappa B_a$ mediate deactivation rate of Cs3 protein | $nM^{-1}$ $h^{-1}$ | [66] |
| $K_{aics3}$ | pAKT mediate deactivation rate of Cs3 protein | $nM^{-K_{n2}}$ $h^{-1}$ | [67] |
| $K_{aic1}$ | Constant controlling the pAKT thresholding effect on Cs3a inhibition | - | [68] |
| $K_{aic2}$ | Constant capturing the saturation effect of pAKT on Cs3a inhibition | $nM^{-1}$ | [69] |
| $K_{n2}$ | Phenomenological exponent to pAKT inhibiting Cs3a | - | [70] |
| $K_{eics3}$ | pERK mediate deactivation rate of Cs3 protein | $nM^{-1}$ $h^{-1}$ | [71] |
| $K_{eic1}$ | Constant controlling the pERK thresholding effect on Cs3a inhibition | - | [72] |
| $K_{eic2}$ | Constant capturing the saturation effect of pERK on Cs3a inhibition | $nM^{-1}$ | [73] |

| | | | |
|---|---|---|---|
| $K_{dbcs3}$ | Deactivation rate of Cs3 protein | $h^{-1}$ | [74] |
| $K_{bcpp}$ | Basal activation rate of CAPP protein | $h^{-1}$ | [75] |
| $K_{acp}$ | $CER_a$ mediated activation of CAPP protein | $nM^{-1}$ $h^{-1}$ | [76] |
| $K_{dcpp}$ | Deactivation rate of CAPP protein | $h^{-1}$ | [77] |
| $S$ | Scaling factor that scaled the basal level of pJNK protein | - | [78] |
| $S1$ | Scaling factor that scaled the basal level of pAKT protein | - | [79] |
| $S2$ | Scaling factor that scaled the basal level of $Cs3_a$ protein | - | [80] |

S2.3: Description of the phenomenological terms used in the model equations

The entire reaction network has been translated into sets of ordinary differential equations (ODEs) (Table S2). While most interactions were represented in the ODEs by mass-action kinetic terms, some were modeled using phenomenological terms that included specific effect of the entities described here.

In Eq. 5, Table S2, the biochemical reaction rate corresponding to inhibition of pJNK by pERK is captured phenomenologically by

$$-\left(\frac{K_{eij}\times pERK\times pJNK}{K_{eij1}+K_{eij2}\times pERK}\right) \qquad [S2.3.1]$$

Note that the rate expression in Eq. S2.3.1 allows introduction of a threshold activation due to pERK mediated inhibition of pJNK.

In Eq. 6, Table S2, we capture the inhibitory effect of TPL on basal NFκB and also activation by TNFR1a by defining the corresponding rates as

$$\frac{(K_{bnf}\times NFkB)}{(1+K_{inh}\times TPL)} \qquad [S2.3.2]$$

and

$$\frac{(K_{tnf}\times NFkB\times TNFR1a)}{(1+K_{inh}\times TPL)}, \qquad [S2.3.3]$$

respectively.

In Eq. 16, Table S2 , the effect of pJNK, pERK and pAKT, respectively on Caspase3 has been introduced phenomenologically by

$$\frac{\left(K_{jacs3}\times Cs3\times pJNK\right)}{\left(K_{jac2}+(n1\times pJNK)\right)} \quad [S2.3.4]$$

$$-\frac{(K_{eics3}\times Cs3a\times pERK)}{\left(K_{eic1}+(K_{eic2}\times pERK)\right)} \quad [S2.3.5]$$

and

$$-\frac{\left(K_{aics3}\times Cs3a\times pAKT^{K_{n2}}\right)}{\left(K_{aic1}+(K_{aic2}\times pAKT^{K_{n2}})\right)} \quad [S2.3.6]$$

Eq. S2.3.4 indicated that pJNK can activate the Caspase3 protein upto a certain threshold level. Eqs S2.3.5 and S2.3.6 ensure that the repression by pERK and pAKT, respectively on Caspase3 is exhibited only above a certain threshold.

The model simulated trajectories scaled with corresponding scaling constant (as shown in Table S4) was used in the objective function (Eq 2, Method M6, Main text) to estimate the parameters based on the experimental training dataset.

### S2.4: Identifiability of the estimated parameters

Parameters were estimated by minimizing the objective function in Eq 2, Method M6, Main text. In order to facilitate the estimation, we considered 2000 fit sequences. For every fit sequence, we initialized the optimization for parameter

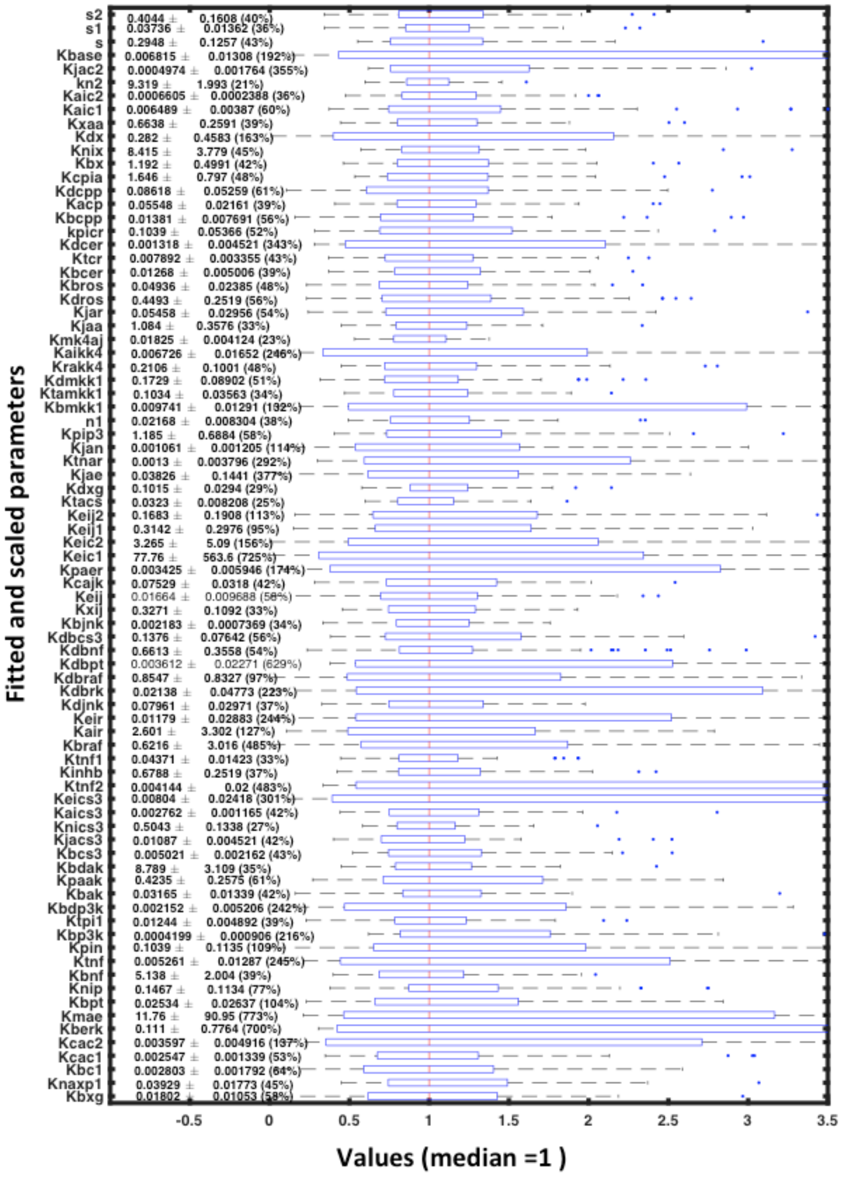


**Figure S4: Boxplot of the estimated kinetic parameters.** Most of the parameters are well constrained and the relative uncertainly in identifying is below 2.5%, which signifies a narrowed distribution of the parameter sets for a multi-experimental fitting.

estimation by randomly generating the starting guess parameter set. The optimization was performed using PottersWheel software (version 4.1.1) (Maiwald et al., 2008) wherein we chose the "Trustregion" method. We chose 3% best fit out of these 2000 sequences that satisfied the adopted statistical fitting criteria in Eq (2), Method M6, Main text. The boxplot in Figure S4 demonstrates the identifiability of these 3% best fit parameter sets.

S2.5: Model trajectories for the best fit parameter set

Out of the 3% best fit parameters, the model trajectories for the one with the lowest $\chi^2$ of 89.21 and Akaike Information Criterion of 488.3 is shown in Figure S5 for all three stimulation conditions along with the corresponding experimentally measured dynamics.

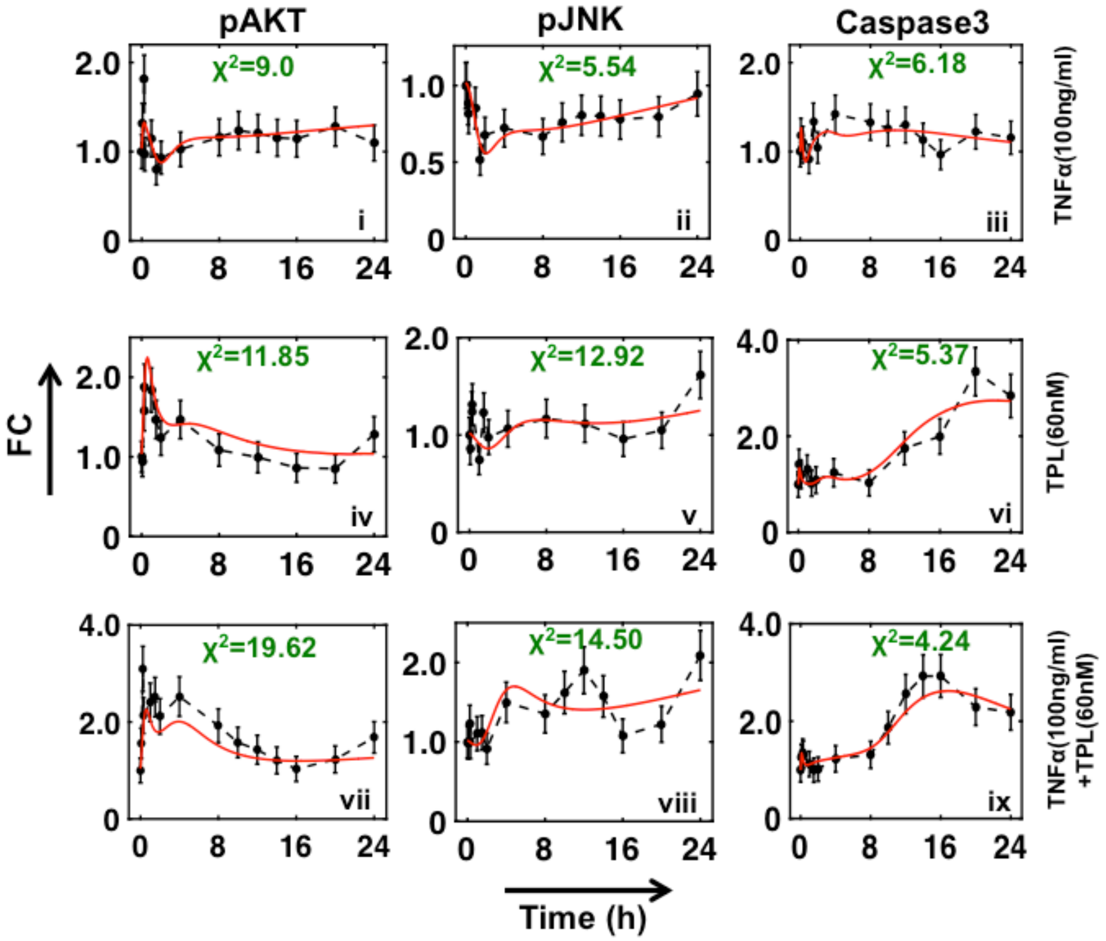


**Figure S5**. **Model trajectories for best fitted parameter set with experimental measurements**. The best-fitted trajectories (red) of ~2000 fits with the experimental FC (circles with appropriate error bars) for pAKT, pJNK and Caspase3 under the three stimulation conditions. The errors are estimated by using a standard error model.

**Text S3: Prediction of independent experimental dynamics by model simulations**

In order to verify the predictive ability of the model, simulations using model equations in Table S2 were performed at two different stimulation conditions, viz., TPL 10 nM and TNFα (100 ng/ml) + TPL (10nM). Experimental measurements on U937 cells under these two stimulation conditions were made (Methods, Main text). The model trajectory predictions and the experimentally measured dynamics are contrasted in Figure S6.

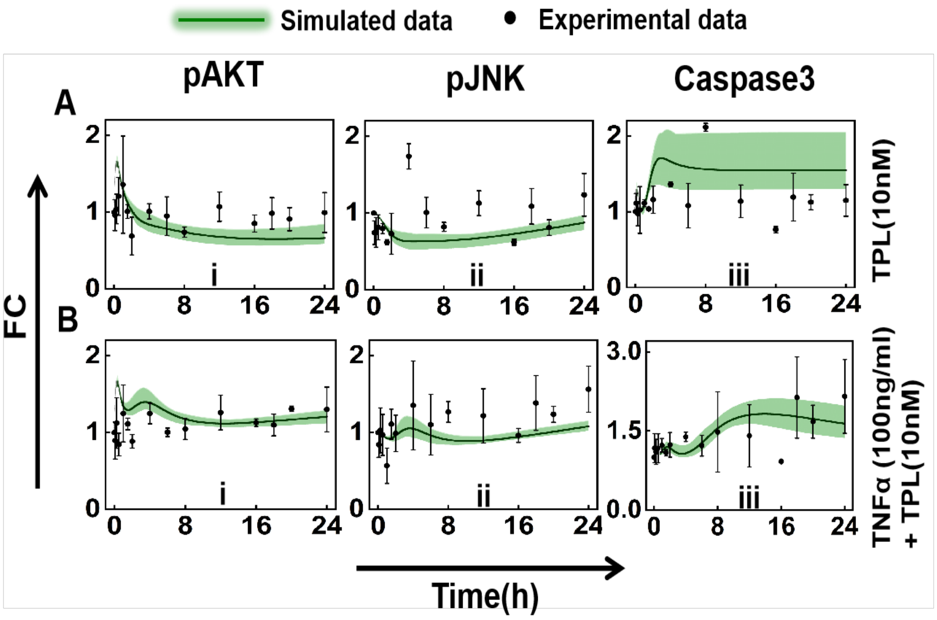


**Figure S6. Comparison of model predicted transients with experimental measurements for TPL-10nM in the presence and absence of TNFα (100ng/ml)**. The green line is the average of best 3% percent parametric values (60 parameter set). Black dots with corresponding error bars (n=3) indicate the experimental measurements. The shaded region represents the minimum and maximum levels that the model transients over 60 parameter sets can achieve.

**Text S4: Quantifying the reaction fluxes influencing the dynamic levels of the three marker proteins.**

The reaction flux analysis was performed to evaluate the contribution of different entities in the TNFα network (Figure S3) in modulating the pAKT, pJNK and Caspase3 dynamics. Flux $J_n$ for all $n = 1,6$ quantifies the rate of each of the 6 terms in the rhs of Eq. 5, Table S2 for pJNK. $A_n$ and$C_n$, respectively capture the fluxes of the terms in the rhs of Eq. 9 (pAKT) and Eq. 16 (Caspase3), Table S2.

**Table S5: Reaction fluxes affecting the pJNK, pAKT and Caspase3 dynamics**

| | |
|---|---|
| $J_n$ indicates fluxes corresponding to different rates in Eq 5, Table S2<br>$\frac{dpJNK}{dt} = (J_1 + J_2 + J_3 + J_4 + J_5 + J_6)$ | |
| $J_1 = (K_{bjnk} \times JNK)$ | [1] |
| $J_2 = (K_{cajk} \times JNK \times C1P_a)$ | [2] |
| $J_3 = -\left(\frac{K_{eij} \times pERK \times pJNK}{K_{eij1} + K_{eij2} \times pERK}\right)$ | [3] |
| $J_4 = -(K_{djnk} \times pJNK)$ | [4] |
| $J_5 = (K_{m4aj} \times JNK \times MEKK_a)$ | [5] |
| $J_6 = -(K_{xij} \times XG_a \times pJNK)$ | [6] |
| $A_n$ indicates fluxes corresponding to different rates in Eq 9, Table S2<br>$\frac{dpAKT}{dt} = (A_1 + A_2 + A_3 + A_4 + A_5 + A_6)$ | |
| $A_1 = (K_{bak} \times AKT)$ | [1] |
| $A_2 = (K_{paak} \times PI3K_a \times AKT)$ | [2] |
| $A_3 = -(K_{bdak} \times pAKT)$ | [3] |
| $A_4 = (K_{jaa} \times pJNK \times AKT)$ | [4] |
| $A_5 = -(K_{cpia} \times CAPP_a \times pAKT)$ | [5] |
| $A_6 = (K_{xaa} \times Bcl2 \times AKT)$ | [6] |
| $C_n$ indicates fluxes corresponding to different rates in Eq. 16, Table S2<br>$\frac{dCs3_a}{dt} = (C_1 + C_2 + C_3 + C_4 + C_5 + C_6 + C_7)$ | |

| | |
|---|---|
| $C_1 = (K_{bcs3} \times Cs3)$ | [1] |
| $C_2 = \left(\frac{K_{jacs3} \times Cs3 \times pJNK}{K_{jac2} + n1 \times pJNK}\right)$ | [2] |
| $C_3 = (K_{tcs} \times Cs3 \times TNFR1_a)$ | [3] |
| $C_4 = -(K_{nics3} \times Cs3_a \times NF\kappa B_a)$ | [4] |
| $C_5 = -\left(\frac{K_{aics3} \times Cs3 \times pAKT^{K_{n2}}}{K_{aic1} + K_{aic2} \times pAKT^{K_{n2}}}\right)$ | [5] |
| $C_6 = -\left(\frac{K_{eics3} \times Cs3_a \times pERK}{K_{eic1} + K_{eic2} \times pERK}\right)$ | [6] |
| $C_7 = -(K_{dbcs3} \times Cs3_a)$ | [7] |

The evolution of the fluxes depicted in Table S5 are shown in Figure S7 for the three stimulation conditions.

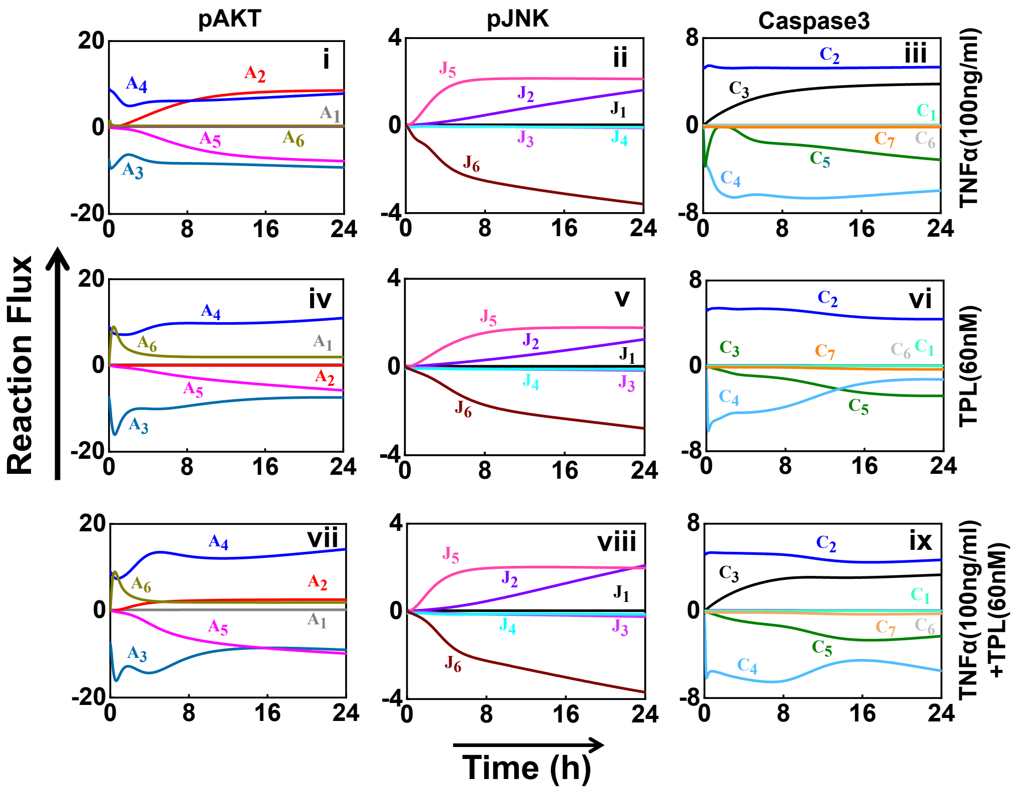


**Figure S7. Flux analysis illustrates the dynamic influence of entities of TNFα network on the marker protein transients.** Three rows correspond to the three stimulation conditions. Expression for various fluxes employed to decipher their dynamic contributions are as given in Table S5.

**Text S5: Model analysis under inhibitory conditions - Wortmannin (Wort) and SP600125 (SP6)**

S5.1: Wort and SP6 Inhibitory model of TNFα network

In order to capture the biological effect of Wort inhibitor on pJNK and pAKT dynamics, we modified Eqs. 5 and 9, Table S2 by introducing its inhibitory action in a phenomenological manner. The modified equations are

$$\frac{dpJNK}{dt} = (K_{bjnk} \times JNK) + (K_{cajk} \times JNK \times C1P_a) - \left(\frac{K_{eij} \times pERK \times pJNK}{K_{eij1} + K_{eij2} \times pERK}\right) - (K_{djnk} \times pJNK) + \left(\frac{K_{m4aj} \times JNK \times MEKK_a}{1 + (K_{kim} \times Wort)}\right) - (K_{xij} \times XG_a \times pJNK) \quad [S5.1.1]$$

and

$$\frac{dpAKT}{dt} = \left((K_{bak} \times AKT) + \left(\frac{K_{paak} \times PI3K_a \times AKT}{1 + (K_{iak} \times Wort)}\right) - (K_{bdak} \times pAKT) + (K_{jaa} \times pJNK \times AKT) - (K_{cpia} \times CAPP_a \times pAKT) + (K_{xaa} \times Bcl2_a \times AKT)\right) \quad [S5.1.2]$$

Similarly, Eqs 5 and 9, Table S2 were modified to incorporate the inhibitory action of SP6 over pJNK and pAKT, respectively as below

$$\frac{dpJNK}{dt} = (K_{bjnk} \times JNK) + (K_{cajk} \times JNK \times C1P_a) - \left(\frac{K_{eij} \times pERK \times pJNK}{K_{eij1} + K_{eij2} \times pERK}\right) - (K_{djnk} \times pJNK) + \left(\frac{K_{m4aj} \times JNK \times MEKK_a}{1 + (K_{kis} \times SP6)}\right) - (K_{xij} \times XG_a \times pJNK) \quad [S5.1.3]$$

and

$$\frac{dpAKT}{dt} = \left((K_{bak} \times AKT) + (K_{paak} \times PI3K_a \times AKT) - (K_{bdak} \times pAKT) + (K_{jaa} \times pJNK \times AKT) - (K_{cpia} \times CAPP_a \times pAKT) + \left(\frac{K_{xaa} \times Bcl2_a \times AKT}{1 + (K_{kib} \times SP6)}\right)\right) \quad [S5.1.4]$$

### S5.2: Model predicted and experimentally measured transients with and without inhibition

A comparison of the inhibitory model transients with that sans inhibition via Wort is in Figure S8(A). This is also juxtaposed with a similar comparison for experimental measurements with and without Wort inhibition conditions. Moreover, these comparisons for the case of with and without SP6 inhibition are in Figure S8(B).

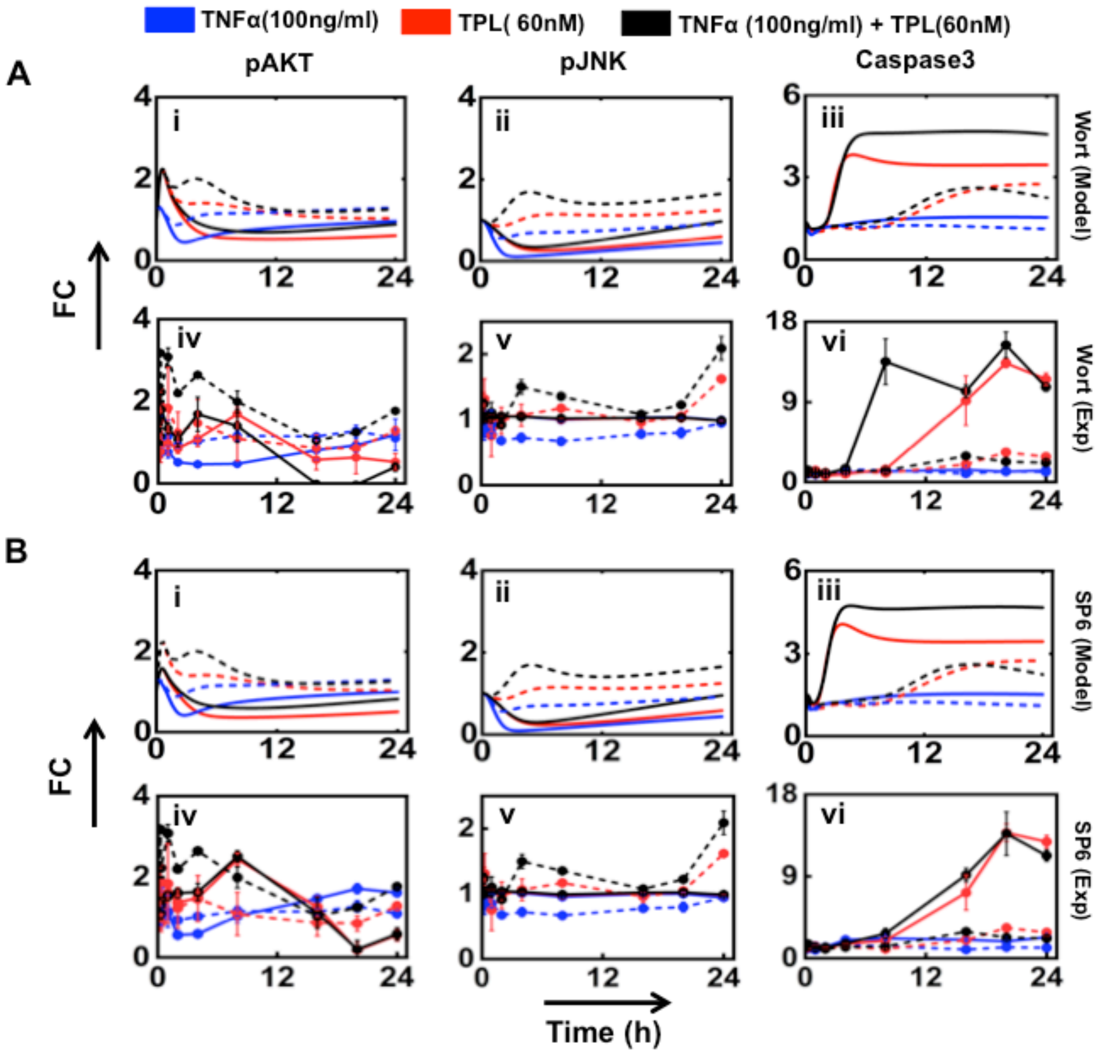


**Figure S8. Comparison of the model and experimental dynamics of the marker proteins under (A) Wort or (B) SP6 inhibitory treatment for all three stimulation conditions.** While solid lines represent inhibitory condition, dashed lines correspond to those without inhibition.

### S5.3: Flux analysis for the Wort and SP6 inhibitory models

In order to perform the flux analysis of the inhibitory models, the modified model specified in section S5.1 along with other equations provided in Table S2 were employed. For the case of Wort inhibition, fluxes J5 and A2 in Table S5 were modified with those rates incorporating its inhibitory action. Similarly, for SP6 inhibition, the inhibitory effects were included in fluxes J5 and A6 in Table S5.

For all three stimulation conditions, we show a comparison of the evolution of fluxes due to important entities corresponding to the three marker proteins in the presence and absence of Wort (Figure S9) and SP6 (Figure S10) inhibition.

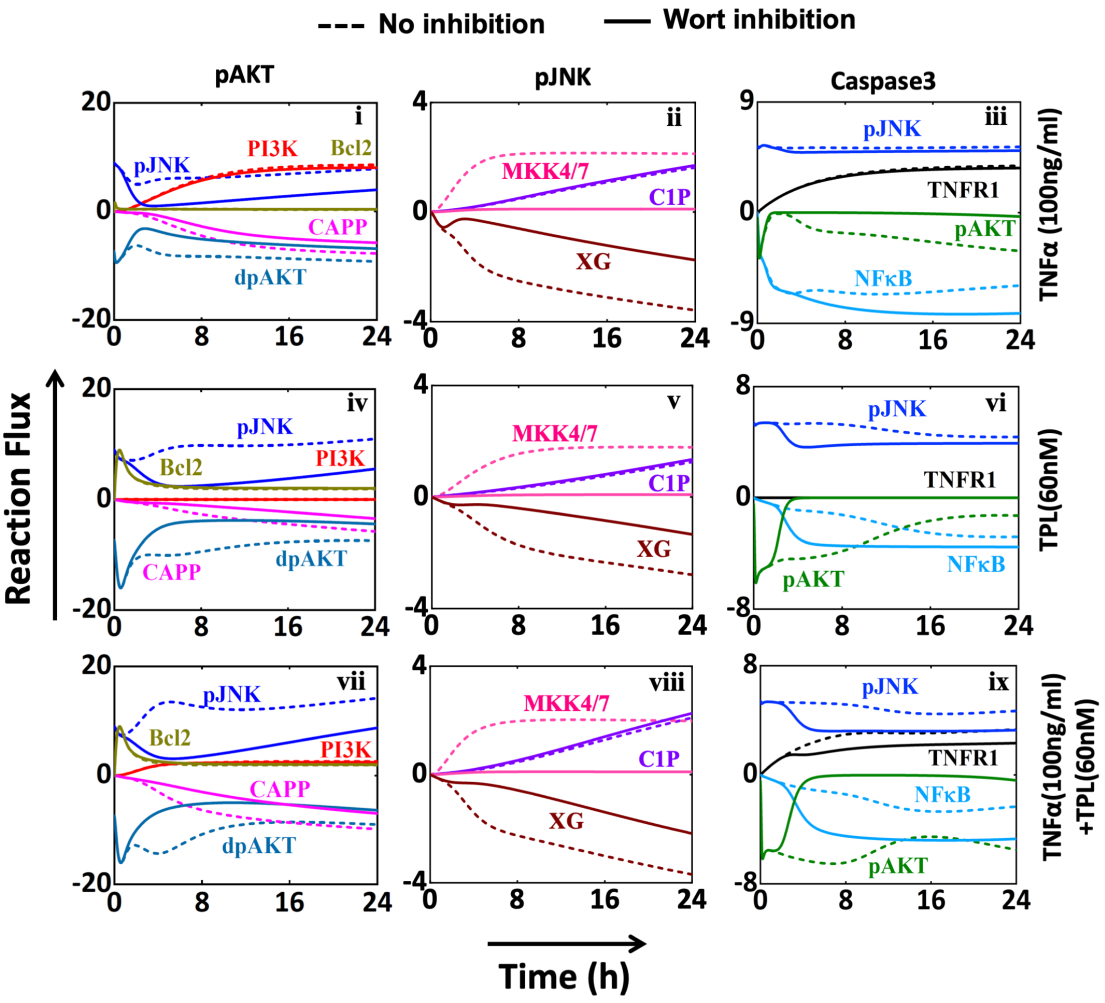


**Figure S9. Evolution of fluxes from important entities controlling the dynamics of pAKT, pJNK and Caspase3 under different experimental conditions in the presence and absence of Wort inhibitor.** Flux analysis of different nodes on controlling of pAKT, pJNK and Caspase3 under different experimental conditions in presence of Wort inhibitor. The contribution from each specific node in the time profile of marker protein has been dissected separately according to Table S5 and then simulated by taking $K_{iak} = 0.001\ nM^{-1}$ and $K_{kim} = 0.02\ nM^{-1}$. The dotted line represents when simulation has been done at *Wort*=0 nM (no inhibitor) and the solid line depicts the trajectories when *Wort*=1000 nM.

The flux analysis for Wort inhibitor (Figure S9) suggests that, in case of pJNK, the inhibition by Wort was more pronounced in suppressing MKK4/7 mediated activation of pJNK. Thus, the effect of the entity MKK4/7 is important under all conditions. Flux analysis suggests that even though Wort directly inhibits pAKT activation via PI3K, it has a moderate influence in regulating pAKT transients. Thus, under the action of Wort inhibitor, pJNK is expected to play a crucial role in regulating cross-talk to modulate Caspase3 transients.

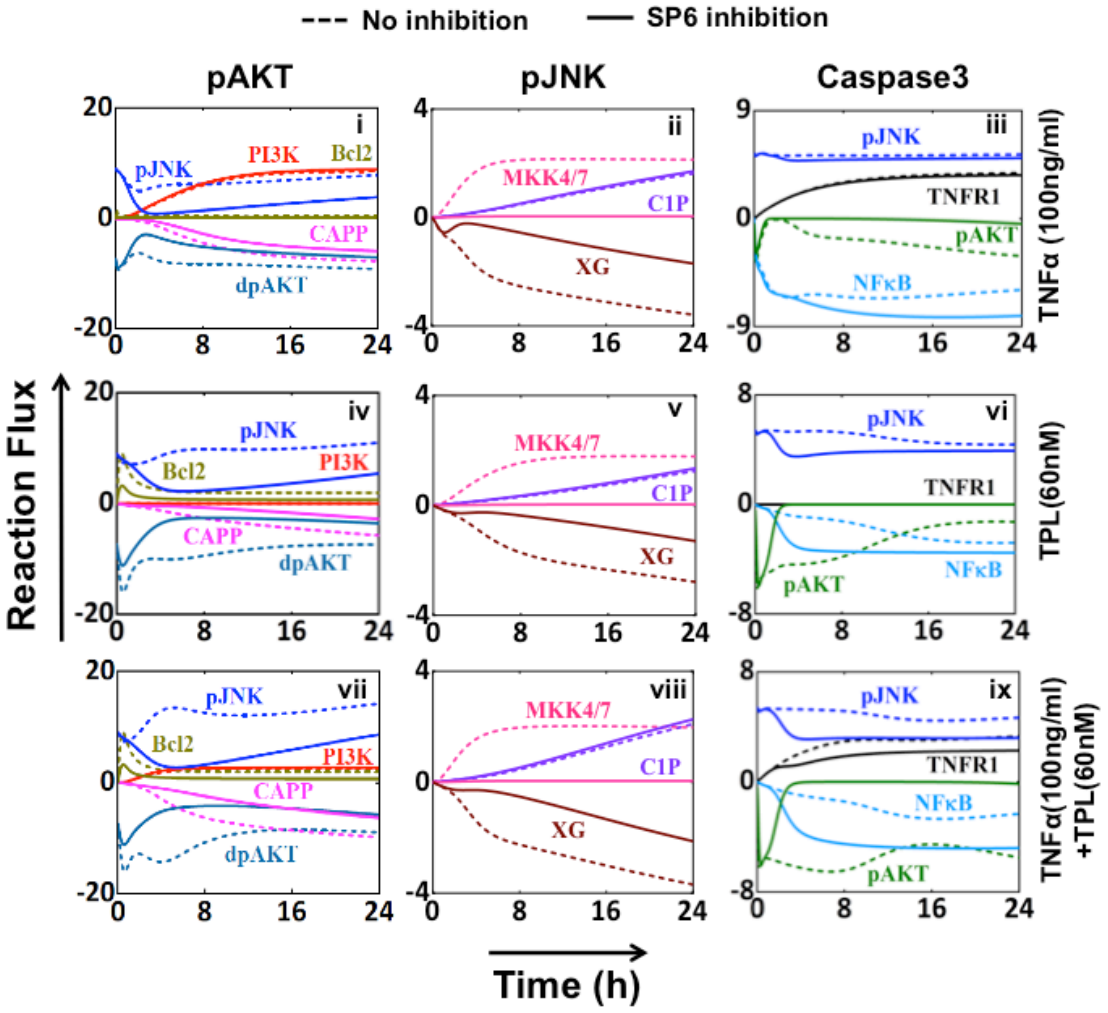


**Figure S10. Evolution of fluxes from important entities controlling the dynamics of pAKT, pJNK and Caspase3 under different experimental conditions in the presence and absence of SP6 inhibitor.** Flux analysis of different nodes on controlling of pAKT, pJNK and Caspase3 under different experimental conditions in presence of SP6 inhibitor. The contribution from each specific node in the time profile of marker protein has been dissected separate according to Table S5 and then simulated by taking as $K_{kib} = 0.0002\ nM^{-1}$ and $K_{kis} = 0.005\ nM^{-1}$. The dotted line represents when simulation has been done at *SP6*=0 *nM* (no inhibitor) and the solid line depicts the trajectories when SP6=$10^4$ *nM*.

Under only TNFα condition, pJNK and PI3K helps to overcome the CAPP facilitated inhibition and control the late activation in the pAKT dynamics in the absence of inhibition. As the retardation effect of Wort on pJNK is comparably less under TNFα stimuli compare to other two conditions (TPL, TNFα+TPL) (Figure S9-(i,iv,vii), blue solid lines), a significant increment has been observed in pAKT dynamics after 12 h when treated with Wort inhibitor (Figure 5A(i), Main text).

The flux study (Figure S9) also reveals that the early and later phase response of Caspase3 dynamics is intriguingly controlled by NFκB, pAKT, and pJNK in a synchronized manner for the three stimulation conditions (Figure 5A(iii)). Under only TNFα condition, as NFκB majorly suppresses Caspase3 activation (Figure S9(iii)), the function of pAKT and pJNK is minimal, resulting in a slight increase in Caspase3 dynamics as compared to no inhibition (Figure 5A(iii), blue line). In absence of Wort, for both only TPL and TNFα+TPL stimulation conditions (Figure S9(vi,ix), dotted line), the early response of Caspase3 is mostly regulated by pAKT and pJNK. Under Wort inhibition, the pAKT negative contribution to Caspase3 significantly reduces after about 5 hours (Figure S9(vi,ix), solid line), resulting in a rapid rise in Caspase3 dynamics at approximately 5 hours (Figure 5A(iii)). Caspase3 maintains a sustained higher level at the later phase as the repression by NFκB has been blocked by the Wort inhibitor (Figure S8(vi,ix), solid line). Thus, the interplay between the pAKT and pJNK mediated inhibition regulates the dynamic response of Caspase3. The flux analysis performed under SP6 inhibitory conditions showed similar responses contributed by different nodes for all conditions as observed under the treatment of Wort (Figure 5B(I,iii) and Figure S10), thereby reproducing similar dynamical response for pAKT, pJNK and Caspase3 in three different experimental conditions.

**Text S6: Semi-quantitative relationship between <AUC$_{casp3}$> and Apoptosis levels**

The relationship between model predicted $\langle AUC_{casp3} \rangle$ and the apoptosis levels in Figure 4B is quantified using a fourth order polynomial function

$$\% Apoptosis = A_0 + A_1\langle AUC_{casp3}\rangle + A_2\langle AUC_{casp3}\rangle^2 + A_3\langle AUC_{casp3}\rangle^3 + A_4\langle AUC_{casp3}\rangle^4 \qquad [S6.1]$$

The coefficients, estimated using Origin software, corresponding to the three stimulation conditions along with $R^2$ are presented in Table S6 below.

**Table S6: Coefficients of the fourth order polynomial (Eq S6.1) under different stimulation conditions.**

| Stimulation condition | Coefficients in Eq. (S6.1) | | | | | $R^2$ |
|---|---|---|---|---|---|---|
| | $A_0$ | $A_1$ | $A_2$ | $A_3$ | $A_4$ | |
| TNFα | -11.78 | 291.10 | -1014.19 | 1371.01 | -610.14 | 99.43 |
| TPL | -49.18 | 783.79 | -2078.08 | 2332.93 | -914.35 | 99.99 |
| TNFα+TPL | -21.72 | 697.59 | -2218.51 | 2831.02 | -1208.87 | 99.98 |

**References**

Allan, L.A., Morrice, N., Brady, S., Magee, G., Pathak, S., and Clarke, P.R. (2003). Inhibition of caspase-9 through phosphorylation at Thr 125 by ERK MAPK. Nat. Cell Biol. *5*, 647–654.

Barkett, M., and Gilmore, T.D. (1999). Control of apoptosis by Rel/NFκB transcription factors. Oncogene *18*, 6910–6924.

Cardone, M.H., Roy, N., Stennicke, H.R., Salvesen, G.S., Franke, T.F., Stanbridge, E., Frisch, S., and Reed, J.C. (1998). Regulation of cell death protease caspase-9 by phosphorylation. Science. *282*, 1318–1321.

Carracedo, A., and Pandolfi, P.P. (2008). The PTEN--PI3K pathway: of feedbacks and cross-talks. Oncogene *27*, 5527–5541.

Deng, Y., Ren, X., Yang, L., Lin, Y., and Wu, X. (2003). A JNK-dependent pathway is required for TNFα-induced apoptosis. Cell *115*, 61–70.

Dhillon, A.S., Hagan, S., Rath, O., and Kolch, W. (2007). MAP kinase signalling pathways in cancer. Oncogene *26*, 3279–3290.

Dobrowsky, R.T., Kamibayashi, C., Mumby, M.C., and Hannun, Y.A. (1993). Ceramide activates heterotrimeric protein phosphatase 2A. J. Biol. Chem. *268*, 15523–15530.

Gangoiti, P., Granado, M.H., Wang, S.W., Kong, J.Y., Steinbrecher, U.P., and Gómez-Muñoz, A. (2008). Ceramide 1-phosphate stimulates macrophage proliferation through activation of the PI3-kinase/PKB, JNK and ERK1/2 pathways. Cell. Signal. *20*, 726–736.

Grummisch, J.A., Jadavji, N.M., and Smith, P.D. (2016). The pleiotropic effects of tissue plasminogen activator in the brain: implications for stroke recovery. Neural Regen. Res. *11*, 1401-1402.

Gustin, J.A., Maehama, T., Dixon, J.E., and Donner, D.B. (2001). The PTEN tumor suppressor protein inhibits tumor necrosis factor-induced nuclear factor-κB activity. J. Biol. Chem. *276*, 27740–27744.

Karin, M. (2002). Lin A.: NFκB at the crossroads of life and death. Nat. Immunol *3*, 221–227.

Lamb, J.A., Ventura, J.-J., Hess, P., Flavell, R.A., and Davis, R.J. (2003). JunD mediates survival signaling by the JNK signal transduction pathway. Mol. Cell *11*, 1479–1489.

Lee, K.Y., Chang, W., Qiu, D., Kao, P.N., and Rosen, G.D. (1999). PG490 (triptolide) cooperates with tumor necrosis factor-α to induce apoptosis in tumor cells. J. Biol. Chem. *274*, 13451–13455.

Mason, C.S., Springer, C.J., Cooper, R.G., Superti-Furga, G., Marshall, C.J., and Marais, R. (1999). Serine and tyrosine phosphorylations cooperate in Raf-1, but not B-Raf activation. EMBO J. *18*, 2137–2148.

Monick, M.M., Powers, L.S., Gross, T.J., Flaherty, D.M., Barrett, C.W., and Hunninghake, G.W. (2006). Active ERK contributes to protein translation by preventing JNK-dependent inhibition of protein phosphatase 1. J. Immunol. *177*, 1636–1645.

Mortenson, M.M., Galante, J.G., Gilad, O., Schlieman, M.G., Virudachalam, S., Kung, H.-J., and Bold, R.J. (2007). BCL-2 functions as an activator of the AKT signaling pathway in pancreatic cancer. J. Cell. Biochem. *102*, 1171–1179.

Ozes, O.N., Akca, H., Gustin, J.A., Mayo, L.D., Pincheira, R., Korgaonkar, C.K., and Donner, D.B. (2005). Tumor Necrosis Factor-α/Receptor Signaling Through the Akt Kinase. In Cell Signaling in Vascular Inflammation, (Springer), pp. 13–22.

Papa, S., Zazzeroni, F., Pham, C.G., Bubici, C., and Franzoso, G. (2004). Linking JNK signaling to NFκB: a key to survival. J. Cell Sci. *117*, 5197–5208.

Park, H.-S., Kim, M.-S., Huh, S.-H., Park, J., Chung, J., Kang, S.S., and Choi, E.-J. (2002). Akt (protein kinase B) negatively regulates SEK1 by means of protein phosphorylation. J. Biol. Chem. *277*, 2573–2578.

Rommel, C., Clarke, B.A., Zimmermann, S., Nuñez, L., Rossman, R., Reid, K., Moelling, K., Yancopoulos, G.D., and Glass, D.J. (1999). Differentiation stage-specific inhibition of the Raf-MEK-ERK pathway by Akt. Science. *286*, 1738–1741.

Rushworth, L.K., Hindley, A.D., O’Neill, E., and Kolch, W. (2006). Regulation and role of Raf-1/B-Raf heterodimerization. Mol. Cell. Biol. *26*, 2262–2272.

Salinas, M., López-Valdaliso, R., Martín, D., Alvarez, A., and Cuadrado, A. (2000). Inhibition of PKB/Akt1 by C2-ceramide involves activation of ceramide-activated protein phosphatase in PC12 cells. Mol. Cell. Neurosci. *15*, 156–169.

Sawada, M., Kiyono, T., Nakashima, S., Shinoda, J., Naganawa, T., Hara, S., Iwama, T., and Sakai, N. (2004). Molecular mechanisms of TNFα-induced ceramide formation in human glioma cells: p53-mediated oxidant stress-dependent and-independent pathways. Cell Death & Differ. *11*, 997–1008.

Shaw, J., and Kirshenbaum, L.A. (2006). Prime time for JNK-mediated Akt reactivation in hypoxia-reoxygenation. Circ Res. *98*, 7-9.

Soga, M., Matsuzawa, A., and Ichijo, H. (2012). Oxidative stress-induced diseases via the ASK1 signaling pathway. Int. J. Cell Biol. *2012*, 439587.

Sohur, U.S., Dixit, M.N., Chen, C.-L., Byrom, M.W., and Kerr, L.D. (1999). Rel/NFκB Represses bcl-2 Transcription in pro-B Lymphocytes. Gene Expr. J. Liver Res. *8*, 219–229.

Vasudevan, K.M., Gurumurthy, S., and Rangnekar, V.M. (2004). Suppression of PTEN expression by NFκB prevents apoptosis. Mol. Cell. Biol. *24*, 1007–1021.

Ventura, J.-J., Cogswell, P., Flavell, R.A., Baldwin, A.S., and Davis, R.J. (2004). JNK potentiates TNF-stimulated necrosis by increasing the production of cytotoxic reactive oxygen species. Genes & Dev. *18*, 2905–2915.

Wajant, H., Pfizenmaier, K., and Scheurich, P. (2003). Tumor necrosis factor signaling. Cell Death & Differ. *10*, 45–65.
